# SRRM2 organizes splicing condensates to regulate alternative splicing

**DOI:** 10.1101/2022.07.04.498628

**Authors:** Shaohai Xu, Soak-Kuan Lai, Donald Yuhui Sim, Warren Shou Leong Ang, Hoi Yeung Li, Xavier Roca

## Abstract

SRRM2 is a nuclear-speckle marker containing multiple disordered domains, whose dysfunction is associated with several human diseases. Using mainly EGFP-SRRM2 knock-in HEK293T cells, we show that SRRM2 forms biomolecular condensates satisfying most hallmarks of liquid-liquid phase separation, including spherical shape, dynamic rearrangement, coalescence, and concentration dependence supported by *in vitro* experiments. Live-cell imaging shows that SRRM2 organizes nuclear speckles along the cell cycle. As bona-fide splicing factor present in spliceosome structures, SRRM2 deficiency induces skipping of cassette exons with short introns and weak splice sites, tending to change large protein domains. In THP-1 myeloid-like cells, SRRM2 depletion compromises cell viability, upregulates differentiation markers, and sensitizes cells to anti-leukemia drugs. SRRM2 induces a *FES* splice isoform that attenuates innate inflammatory responses, and *MUC1* isoforms that undergo shedding with oncogenic properties. We conclude that SRRM2 acts as a scaffold to organize nuclear speckles, regulating alternative splicing in innate immunity and cell homeostasis.

## INTRODUCTION

Proteins with intrinsically disordered regions (IDRs), often together with RNA, can form biomolecular condensates via phase separation to form liquid-like membraneless organelles (1), which are rapidly and reversibly organized for the regulation of biochemical processes within cells (2, 3). Nuclear speckles are a type of membraneless organelles in nucleus, acting as reservoirs of splicing factors, although the cellular function of the nuclear speckles is not clear (3, 4).

Previous studies indicated that nuclear speckles are composed of two layers, with SRSF2 and SON in the center and other splicing factors and RNAs like MALAT1 long-noncoding RNA or U1/U2 small nuclear RNAs (snRNAs) in the periphery (5). However, only recently the SRSF2 antibody (SC35) was found to cross-react with SRRM2, suggesting that the key component of speckles is indeed SRRM2 (6). Nuclear speckles dissipate after nuclear envelope breakdown and entry into metaphase, and speckle proteins form distinct droplets named Mitotic Interchromatin Granules (MIGs) (7). Although nuclear speckles appear similar to other membraneless organelles demonstrated to undergo phase separation such as nucleoli, the biophysical mechanism underlying nuclear-speckle formation has not been formally demonstrated (8). This knowledge gap is important, considering the recent controversy over studies invoking phase separation without direct and conclusive evidence (9).

Certain splicing factors possess properties related to phase separation to regulate alternative splicing independently of RNA recognition motifs (RRM). The Rbfox1 C-terminal tyrosine-rich domain promotes aggregation, nuclear-speckle localization, and splicing activation (10). Furthermore, glycine/tyrosine-rich low-complexity regions in mammalian hnRNPs remodel protein-interaction networks and affect the formation of higher-order assemblies on target pre-mRNA, causing exon skipping (11). RNA facilitates phase separation of splicing factors such as hnRNPA1 (12). Several IDR-containing proteins inside nuclear speckles have an inherent property to phase separate, including SRSF1, SRSF2 and U2AF2, and they form condensates incorporating hyperphosphorylated RNA polymerase II (Pol II) C-terminal domain for splicing regulation (13). Nevertheless, the properties and functions of most IDRs in splicing factors remain uncharacterized. Here we provide evidence for SRRM2 as key organizer of speckle formation, likely via liquid-liquid phase separation.

Like its much smaller budding yeast orthologue Cwc21, human SRRM2 (SRm300) is an RS-repeat containing protein found in spliceosomal structures by cryo-electron microscopy (EM) (14), validating prior biochemical studies (15, 16). The structured N-terminus of SRRM2/Cwc21 sits inside pre-catalytic complexes and contacts the end of the 5’ exon throughout the trans-esterification reactions, while the large SRRM2 IDRs could be attributed to a nearby lobe of weak EM density (14). By virtue of its spliceosomal location and interaction with CWC22, SRRM2 might help deposit the Exon Junction Complex (EJC) onto mature mRNAs after splicing. SRRM2 and its smaller and also IDR-rich SRRM1 homologue form a splicing co-activator complex that supports both constitutive and ESE-dependent splicing in cell-free extracts (17). Certain diseases feature SRRM2 sequence and expression abnormalities alongside altered splicing patterns (18, 19), suggesting that SRRM2 regulates alternative splicing independently of RRM. DYRK3 inhibition led to the formation of aberrant SRRM2 hybrid condensates with proteins from various membraneless organelles (7), and SRRM2 forms cytosolic condensates if mislocalized (20). Among DYRK3-specific interactors, SRRM2 was the chief splicing-speckle protein with multiple phospho-sites peaking at mitosis and downregulated upon DYRK3 inhibition (7,21,22). Additionally, human SRRM2 depletion disrupts the localization of splicing factors (like PQBP1) and RNAs (U1 snRNA and poly(A)^+^ RNA) to nuclear speckles (20, 23). SRRM2 appeared to be important for immune cells, as it is strongly downregulated during differentiation of human primary macrophages (24), its mutation increases exon skipping in leukocytes (18), and its depletion alters cytokine secretion in immune cells (25). All this embodies the importance of elucidating the SRRM2 properties in relation to phase separation, and their influence in alternative splicing in immune cells.

## MATERIALS AND METHODS

### Cell Culture

We cultured HEK293T (ATCC, CRL-11268) in Dulbecco’s Modified Eagle’s Medium (DMEM) (Hyclone, SH30022.01) and THP-1 (ATCC, TIB-202) in RPMI-1640 (ATCC, 30-2001), both supplemented with 10% FBS (Gibco, LS26140079), 100 U/ml Penicillin and 100 μg/ml Streptomycin at 37 °C, and 5% CO_2_. We differentiated Macrophage-like THP-1 cells following the procedures as previously described (24). For confocal imaging, we grew cells on glass coverslips (CellPath, SAH-2222-03A), coated with poly-L-lysine (Sigma Aldrich, P4707). We show all key resource information in Key Resources Table.

### Knock-in cell line generation

We used the CRISPR/Cas9 system to generate HEK293T cell lines with SRRM2 endogenously tagged with monomeric enhanced GFP (mEGFP), and SRSF2 endogenously tagged with mCherry. We cloned the repair templates into a pUC19 vector containing fluorescent protein (EGFP to C-terminus of SRRM2) or mCherry to N-terminus of SRSF2), a GST linker (G(GST)_4_GG for SRRM2 and G(GST)_3_GG for SRSF2), and 800 bp flanking homology arms for recombination. We co-transfected HEK293T cells with the repair templates and pX330 vector containing the corresponding sgRNAs, and selected clones by serial dilution followed by genomic genotyping. We show the sequences of sgRNAs and genotyping primers in Supplementary Table 1. We chose homozygous clones for the experiments.

### Plasmids and cloning procedures

We amplified Complementary DNAs (cDNAs) of genes by PCR using PrimeSTAR Max DNA polymerase (TAKARA Bio), and cloned the PCR products into overexpression vectors EGFP-N1 (Addgene) or pLenti6 (Addgene). We performed mutagenesis using KAPA HiFi polymerase (KAPA Biosystems). For splicing minigenes, we amplified FES exons 9 to 11 with intervening sequences and SH3BP2 exons 9 to 11 with intervening sequences from human genomic DNA (Promega), and cloned the products into pcDNA3.1(+) plasmid (Addgene). For knockdown plasmids, we inserted shRNA sequences into pLKO.1 (Addgene) or shmirRNA (Addgene) by DNA oligonucleotide annealing and ligation. We used shmirRNA for reducing the off-target effect of shRNA. We used pET24b (+) for constructing protein expression plasmids in *E. coli*. We show KSR and UPR_3_ protein sequence in Supplementary Table 1.

### Lentiviral Transduction

We co-transfected HEK293T cells with overexpression plasmids or shRNA viral plasmids with lentiviral packaging vectors using X-tremeGENE 9 (Sigma-Aldrich), and we collected viruses at 48 h post-transfection. We infected THP-1 cells with overexpression or shRNA lentiviruses and induced them to differentiate with 20 ng/ml PMA 72 h post-infection. We listed the oligonucleotides used in this study in Supplementary Table 1.

### Semi-quantitative RT-PCR and qRT-PCR

We extracted total RNA using QIAGEN kit (QIAGEN, 74106). We generated cDNA with random hexamers for qRT-PCR) or oligo(dT) primers for RT-PCR using M-MLV (NEB). We carried out semi-quantitative RT-PCR using DreamTaq (Thermo Fisher), examined the PCR products by agarose gel electrophoresis, and imaged the gels using the GeneSys programme on a T:Genius gel doc (SynGene, USA). We then analysed the intensity of the bands using ImageJ to calculate the PSI values, and we then compared them to the PSI values of the corresponding DASEs obtained from RNA-seq, using Pearson’s correlation coefficient.

*Pearson^’^s correlation coefficient* 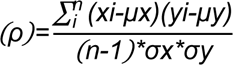 *μ=mean σ=standard deviation* We performed SYBR Green reagent-based qRT-PCR on Bio-Rad CFX Real-time PCR System. We used Actin as an internal control for normalization. We listed the primers and length of products in Supplementary Table 2.

### Western Blotting

We prepared whole cell lysates in ice-cold cell lysis buffer (Sigma-Aldrich). We separated Protein samples on a TGX gel (Bio-Rad), and blotted them onto PVDF membranes (Millipore) at 110 V for 1.5 h. After blocking in 0.02 M Tris-Base, pH 7.6, 2% milk powder, 0.137 M NaCl, 0.1% Tween 20, we incubated the membranes with a primary antibody at 4 °C overnight, then incubated with secondary horseradish peroxidase-conjugated anti-mouse antibody, followed by immunodetection with SuperSignal West Pico Chemiluminescent Substrate (Thermo Fisher). We recorded images with a ChemDoc MP Image System (Bio-Rad).

### Immunofluorescence

We grew HEK293T and differentiated THP-1 cells on coverslips, and prepared suspension THP-1 cells by cytospin. We fixed the cells with 4% PFA for 5 min and permeabilized them with 0.5% Triton X-100 for 5 min. We blocked the sample with 1% FBS in PBS, incubated with primary antibodies at 4 °C overnight, and then incubated with Alexa-Fluor 647 labelled secondary antibody (Biolegend) for 1 h at room temperature. We stained nuclei using DAPI (Life Technologies).

### Flow cytometry

We resuspended cells in staining buffer (PBS with 10% FBS) with 1×10^6^ cells/ml, and blocked the nonspecific staining by anti-CD16/32 for 10 min on ice. We then stained the cells with monoclonal antibodies (mAbs) at 4 °C for 30 min, and washed them with PBS. We measured marker expression on the BD LSRFortessa (BD) and analyzed with FlowJo software. We purchased fluorochrome-conjugated mAbs and the corresponding isotype controls from Biolegend, BD Biosciences, or eBioscience, as indicated in Supplementary Methods file.

### 4D time-lapse imaging and line scan analysis

We grew cells on glass-bottomed 35 mm µ-Dish (ibidi, 81156). We used Hoechst 33342 (0.5 μg/ml, Thermo Fisher Scientific) to stain DNA 1h before imaging, and changed the medium to Leibovitz’s L-15 media (no phenol red) (Gibco, 21083-027). We maintained all cells at 37 °C under 5% CO_2_. We carried out 4D time-lapse imaging on a Zeiss LSM980 with Airyscan 2 confocal microscope (Carl Zeiss) using a Plan-Apochromat 63x/1.40 oil objective. We imaged cells with Z stacks (0.2 to 0.4 μm apart, 50 to 70 stacks) which we acquired at each time point. We performed time-lapse imaging with a time interval of 5 to 20 min. We acquired images using the multiplex SR-4Y mode, and subsequently processed them using the Zen 2 (blue edition) software (Carl Zeiss). We carried out line scan profile analysis (and number/size of condensates) using ImageJ. We drew line scan lines using the “line tool” from Image J through the region of interest. We plotted the pixel intensity along the line and normalized it to a maximum of 1. We identified spots with threshold by IMARIS software.

### FRAP in live cells

We performed live cell Fluorescence Recovery After Photobleaching (FRAP) on a Zeiss LSM 710 confocal microscope equipped with a 1.46 numerical aperture (NA) Plan-Apo ×100 oil-immersion lens (Carl Zeiss, Germany). We performed photobleaching using 75% laser power (488 nm wavelength of an argon laser). We imaged the photobleached region every 500 ms for 75 cycles, with laser power attenuated to 1.5% intensity. We measured the fluorescence intensity at the bleached spot, a control unbleached spot, and background. We analyzed the data using Zen 2.3 SP1 (black) software (Carl Zeiss).

### Drug treatments

For transcription inhibition, we incubated THP-1 cells with 50 μM DRB or 1 μM CPT (camptothecin) (Sigma) for 7 h. To induce proinflammatory cytokines in human macrophage-like THP-1 cells, we treated these cells with LPS (1 ng/ml) for 6 h. For AML drug treatment, we incubated either control THP-1 or SRRM2 knockdown THP-1 cells with different concentrations of CPT or AZA (Sigma) for 24 h. We employed the celltiter-glo luminescent cell viability assay kit (Promega) and luminescence Tecan infinite M200Pro plate reader (TECAN) to assay the cell viability.

### Protein purification

We produced SRRM2-IDRs in *E. coli* strain Rosetta (DE3) cells. We grew cells in Lysogeny broth medium to OD600 0.8 at 37 °C, then we added isopropyl β-D-1-thiogalactopyranoside (1 mM) to induce protein expression from pET24b (+) for 3 h. We lysed cells in high-salt lysis buffer (20 mM Tris-HCl, pH 8.0, 500 mM NaCl, and 10 mM imidazole) in the presence of lysozyme (0.3 mg/ml) and 2 mM phenylmethanesulfonyl fluoride (PMSF), then treated samples with RNase A (5 U/ml) and TURBO DNase (1 U/ml, Thermo fisher) for 30 min, and sonicated them for 15 min. We loaded the soluble fraction on an IMAC (HisTrapTM HP 5 ml, GE Healthcare) column, and washed it with wash buffer (20 mM Tris-HCl, pH 8.0, 250 mM NaCl, and 10 mM imidazole). We eluted bound proteins using an imidazole gradient and subjected them to size-exclusion chromatography (Superdex 200 16/600, GE Healthcare) using 20 mM Tris-HCl (pH 8.0) and 250 mM NaCl as eluent.

### *In vitro* droplet assay

We concentrated recombinant EGFP fusion proteins using Amicon Ultra centrifugal filters (10K MWCO, Millipore) to an appropriate protein concentration in 250 mM NaCl salt buffer. We added recombinant proteins to solutions at varying concentrations with 75 to 300 mM final salt buffer (in 20 mM Tris-HCl pH 7.5) with or without 5% PEG-8000 as crowding agent or RNA (extracted from HEK293T cells and purified as we did for RNA-seq to final concentration of 100 ng/μl). We mixed the protein solution and added it onto a coverslip for imaging. We took images in 2 mins for normal condensate formation experiments with a Nikon microscope with 100X objective.

### RNA-sequencing and data analysis

We isolated Total RNA using RNeasy Mini Kit (Qiagen), followed by DNase treatment (TURBO DNA-free Kit, Ambion) and RNA purification (RNA Clean and Concentrator, Zymo Research). We prepared cDNA libraries using the TruSeq Stranded mRNA Library Prep Kit (Illumina) as described (24, 26). We used HiSeq 2500 System Rapid mode (Illumina) for RNA-sequencing, and confirmed the quality using FastQC (Simon Andrews, Babraham Bioinformatics). We aligned the paired-end reads to human genome (UCSC-hg19) using Tophat2 (27) with Bowtie2 (28). We employed rMATS to analyze alternative splicing events (29). We used the default parameters for each tool unless otherwise stated.

## RESULTS

### Nuclear-speckle condensation via SRRM2 in cell cycle

To clarify how SRRM2 affects the integrity of nuclear speckles, we used shRNAs to knock down the core components SRRM2 and SON in HEK293T cells by shRNAs concomitant with fluorescent labeling of transfected cells (Figure S1A). Then we visualized SRRM2, SON, or SRRM1 as nuclear-speckle makers. We found that the cells with a single knockdown of either SRRM2 (only green) or SON (only red) did not disrupt the foci with SRRM1 or the other factor. In turn, cells with double knockdown of SRRM2 and SON (both green and red) dispersed SRRM1, which is indicative of nuclear-speckle disassembly (Figure 1A). This result is consistent with a recent study that used siRNA knockdown (6), and indicates that both SRRM2 and SON contribute to nuclear-speckle integrity. Since SRRM2 alone is essential for the localization of some splicing factors and especially U1 snRNA or poly(A)+ RNAs to nuclear speckles (20, 23), SRRM2 appears more important for nuclear-speckle integrity.

**Figure 1.**
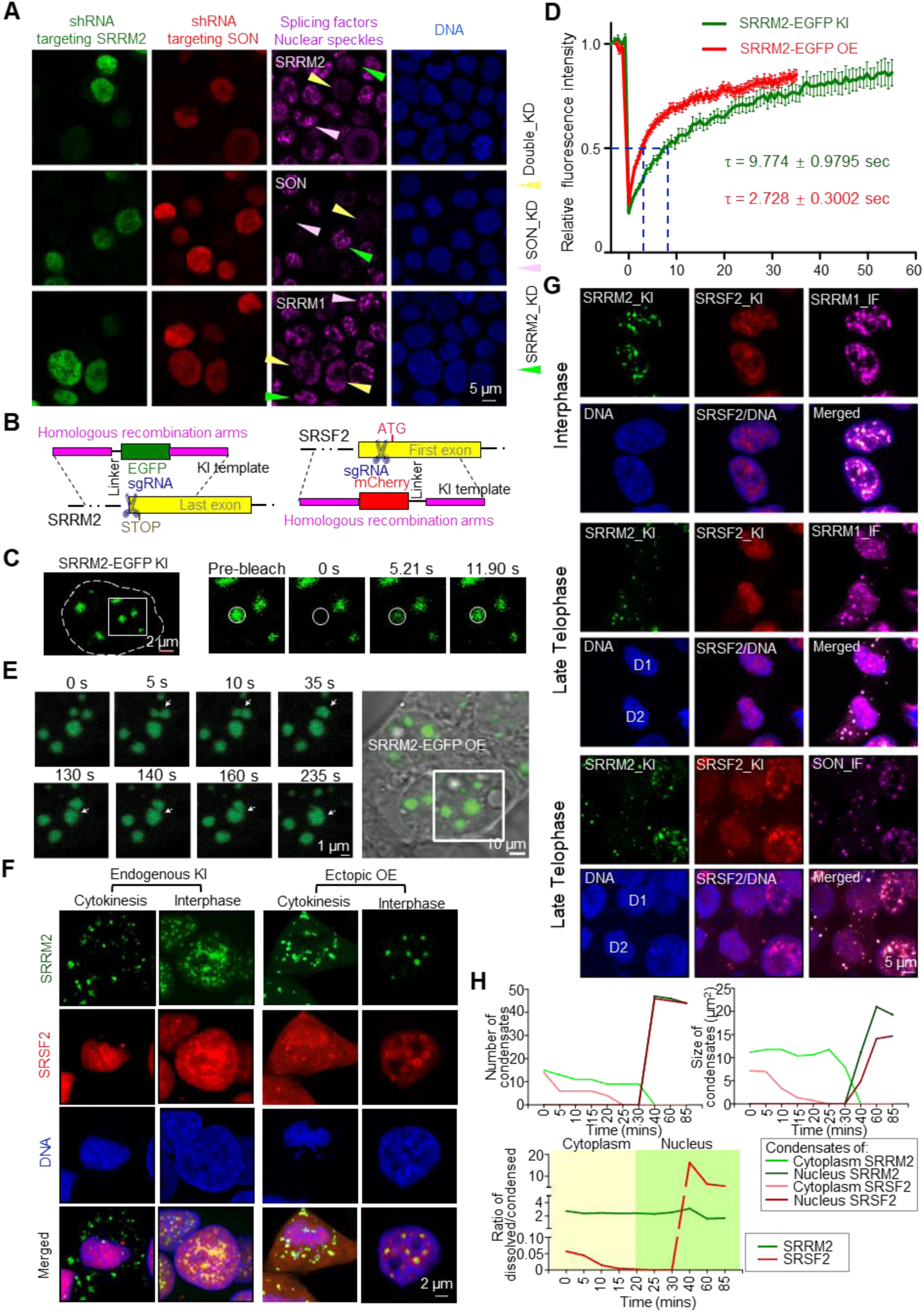
SRRM2 is responsible for organizing nuclear speckles. (**A**) Representative images of SRRM2, SON, or SRRM1 immunofluorescence staining upon single knockdown of SRRM2 (green cells and arrows), SON (red cells and pink arrows), or SRRM2/SON double knockdown (green and red, yellow arrows) in HEK293T cells (experimental design in Figure S1A). (**B**) Schematic of the experimental design of SRRM2-EGFP and SRSF2-mCherry knock-ins by CRISPR/Cas9 system. (**C**) Representative images of FRAP experiments on knock-in EGFP-SRRM2 in HEK293T cells. (**D**) Fluorescence recovery curve of the FRAP experiment of either knock-in or overexpressed EGFP-SRRM2 with relative fluorescence intensity of EGFP-SRRM2 plotted against time. (**E**) Live-cell images of two fusion events (arrowheads) among three SRRM2 condensates. Two condensates fused at 5 s, and the formed liquid condensate fused with another one at 140 s. See Video 1. (**F**) Live-cell imaging 3D reconstruction showing the distribution of SRRM2-EGFP and SRSF2-mCherry at cytokinesis or interphase stage, in either double knock-in (KI) cells or ectopic overexpression (OE) through co-transfection, all in HEK293T. (**G**) Immunofluorescence staining on fixed double knock-in cells showing the distribution of SRRM1 and SON in late telophase and interphase. (**H**) Number and size of condensates of SRRM2 or SRSF2 in cytoplasm or nucleus, and ratio of dissolved/condensed SRRM2/SRSF2 in cytoplasm or nucleus at different cell cycle stages by 4D live-cell imaging (Figure S1F and Video 2). Images are representative of at least three independent experiments.

To investigate the condensation properties of SRRM2, we generated a SRRM2-EGFP knock-in cell line in HEK293T by CRISPR/Cas9 (Figure 1B and S1B). With these cells, we performed Fluorescence Recovery After Photobleaching (FRAP) and time-lapse analysis of SRRM2 condensates in the nucleus (Figure 1C). Knock-in SRRM2-EGFP localized to micron-sized spherical condensates that recovered with a half-time of ∼9.7 s (Figure 1D), which is slightly slower than ectopically overexpressed SRRM2-EGFP (Figure S1C). When SRRM2 condensates came in contact, we observed rapid fusion followed by relaxation to a spherical shape (Figure 1E). Both coalescence and dynamic rearrangement are hallmarks of phase-separated condensates.

We hypothesized that the splicing factors with potential for phase separation behave differently along the cell cycle, compared to other nuclear-speckle splicing factors and components (Figure S1D). To test this, we generated SRRM2-EGFP and SRSF2-mCherry double knock-in HEK293T cells. By live-cell 3D-imaging in these cells, we observed that the dynamics of SRRM2 and SRSF2 condensates were highly distinct from later telophase to interphase (Figure 1F). With both knock-in and ectopic overexpression, SRRM2 and SRSF2 colocalized in nuclear speckles as expected. Importantly during cytokinesis, SRSF2 accumulated in chromatin regions that are clearly separate from MIGs where SRRM2 condensed. Upon overexpression, SRRM2 localized to the core while SRSF2 localized to the external layer of SRRM2 condensates, indicating a multi-layered organization of nuclear speckles (Figure 1F). By immunofluorescence staining of endogenous SRRM1 and SON in SRRM2-SRSF2 double knock-in cells, we showed that SRRM1 localized with both SRRM2 and chromatin/SRSF2 while SON localized entirely with SRRM2 in late telophase (DKC1 staining showing nucleolar condensation) (Figure 1G and Figure S1E). These results suggest that the different cell cycle dynamics of the nuclear-speckle proteins reflect a distinct role in condensate formation.

To investigate how nuclear speckles form after mitosis, we further tracked the dynamics of these splicing-speckle proteins by live-cell 3D imaging. From telophase, condensed SRSF2 gradually dissolved, and dispersed SRSF2 located to chromatin, at the time that DKC1 already formed condensates within chromatin region (Figure S1E and S1F). In turn, condensed SRRM2 was retained in the cytoplasm, even when the nuclear envelope appeared and all SRSF2 was imported into the nucleus. Later, cytoplasmic SRRM2 condensates slowly dissolved and started to enter the nucleus (Figure S1F). Speckles only condensed when SRRM2 entered the nucleus, which triggered an increase in the ratio between condensed and dissolved SRSF2. The intensity and size of splicing condensates increased along with SRRM2 intensity inside the nucleus (Figure 1H and S1G). Overall, these results strongly suggest that SRRM2-mediated nuclear condensation is essential for nuclear-speckle formation.

### SRRM2-Intrinsically Disordered Regions drive condensate formation

Our evidence suggests that phase separation may be the mechanism by which SRRM2 forms condensates. To predict the sequences that are responsible for condensate formation likely via phase separation, we performed disordered tendency, FOLD index, and NCPR analysis of the human SRRM2 protein (30–32). We found a similar domain structure for SRRM2 as previously reported (33). In particular, the conserved lysine/serine/arginine-rich (KSR) region, as well as the undeca- and dodeca-amino acid repeat regions (UPR and DPR respectively) (Figure S2A) are highly disordered and possess a net positive charge (Figure 2A). Based on these predicted IDRs, domain mapping of SRRM2-truncated GFP fusion genes highlighted few different properties of SRRM2 IDRs (Figure S2B and S2C). First, the SRRM2-mutant named ΔNCK, without the conserved NCK domains (<u>N</u>-terminus, <u>c</u>wf21 domain, and <u>KSR</u>) formed small granules. Second, mutants containing the DPR and nearby RS-rich region formed foci likely corresponding to aggregates, which is consistent with a recent report showing that the C-terminus of SRRM2 forms aggregates with tau in the cytoplasm (34). Third, mutants with NCK plus DPR formed condensates. Additionally, the double UPR and DPR deletion mutant lost the condensate-formation properties, as most of this protein was dispersed inside the nucleus (Figure 2B). To rule out spurious effects of overexpression, we found that ΔUPRΔDPR did not form condensates and instead dispersed in the nucleus, even with an expression level higher than the full-length protein in SRRM2 knockdown cells (Figure 2C). Moreover, the NCK fragment formed condensates in the cytosol when fused to a Nuclear Export Signal (NES), indicating that NCK induces condensate formation independently of proper nuclear localization (Figure 2D). We conclude that NCK, UPR, and DPR are responsible for the formation of SRRM2 condensates in cells.

**Figure 2.**
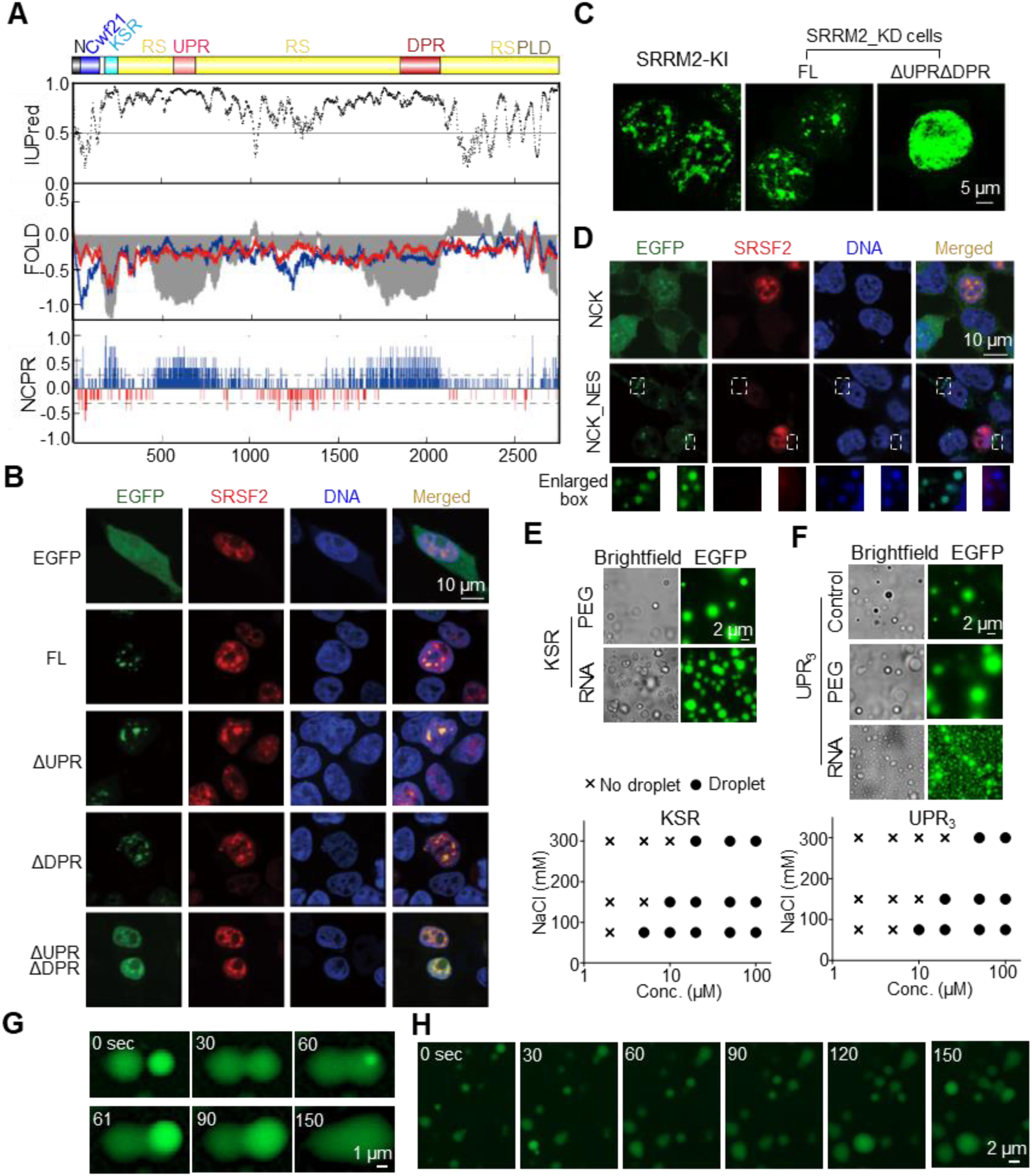
SRRM2-IDRs are responsible for condensate formation. (**A**) Predicted IDRs in human SRRM2. IUPred for disordered tendency; FOLD: intrinsic disorder prediction by PLAAC (blue) and PAPA (red); and NCPR: net charge per residue; RS: RS-rich region; PLD: prion-like region (PLAAC). (**B**) Immunofluorescence images of HEK293T cells co-transfected with EGFP-SRRM2 with mCherry-SRSF2 and single deletion of UPR, DPR or UPR/DPR double deletion. FL: Full-length SRRM2. (**C**) Images showing the localization of either FL SRRM2 or ΔUPRΔDPR in SRRM2 knockdown HEK293T cells. SRRM2-EGFP knock-in cell as a reference for showing condensates at a relatively similar expression level (images taken and displayed under same parameters). (**D**) Images showing control-NCK and NES (Nuclear Export Signal)-NCK with EGFP tag expressed in HEK293T cells. We labeled DNA in fixed cells with DAPI. Enlarged boxes show DAPI accumulated on NCK condensates in the cytoplasm. (**E and F**) *In vitro* phase-separation experiments showing protein condensates at varying NaCl (75, 150, or 300 mM) and peptide KSR (E) and UPR_3_ (F) (2, 5, 10, 20, 50, 100 μM) concentrations under 5% PEG. Phase diagram plots the presence (circles) or absence (crosses) of droplets. We show a representative image of 20 μM under 150 mM NaCl (control), protein droplet formation of KSR after adding RNA (final concentration 100 ng/μl) without PEG, and UPR_3_ droplet formation with PEG (5% final concentration) or RNA (100 ng/μl final concentration) as representative pictures. (**G**) Representative image of two fusion events of KSR-EGFP droplets formed *in vitro* at concentration of 20 μM under 150 mM NaCl and 5% PEG. Droplet fusion started at 30 sec and 61 sec. (**H**) Time-lapse of UPR_3_ droplets at concentration of 20 μM under 150 mM NaCl. We observed several fusion events at each time point. See Video 3.

Next we sought to elucidate whether the KSR or low-complexity repeat UPR/DPR IDRs fused to EGFP form liquid-like condensates *in vitro* under physiological salt conditions (100–150 mM NaCl) (Figure S2D). We did not observe droplet formation of KSR even at a concentration as high as 100 μM under low salt concentration (50 mM NaCl), which promotes phase separation of many known IDRs (11, 12). In the presence of 5% polyethylene glycol (PEG), the KSR protein phase separated from a concentration as low as 5 μM, indicating the necessity of molecular crowding (Figure 2E). UPR_3_ (3 repeats of the eleven/undeca-amino-acid sequence) phase separated without crowding agent, yet 5% PEG promoted larger condensate formation, indicating that its droplet formation propensity increased with higher molecular crowding (Figure 2F). As expected, KSR and UPR_3_ phase diagrams revealed that protein concentration affected droplet formation, fulfilling another hallmark of phase separation. Lowering the NaCl concentration led to condensate formation at lower protein concentrations, suggesting that electrostatic interactions contribute to SRRM2 droplets. These condensates were liquid-phase separated as they were spherical, concentration dependent and capable of fusion (Figure 2G and 2H). Importantly, total RNA promoted condensate formation of KSR without crowding agent, and the UPR_3_ droplets became smaller yet with higher intensity upon adding RNA. Moreover, both NCK-NES condensates in the cytosol, and the nuclear UPR or DPR unexpectedly accumulated DAPI staining, unveiling the intrinsic ability of these IDRs to recruit nucleic acids (Figure 2D and S2E). We conclude that SRRM2-IDRs are responsible for biomolecular condensate formation likely via phase separation, and that the condensates coacervate with RNA.

### SRRM2 is associated with myeloid leukemia by promoting cell survival and repressing differentiation

SRRM2 was associated with several human pathologies such as neurodegenerative diseases and cancers (18, 20). We obtained the expression levels of SRRM2 in different types of cancers in TCGA database. SRRM2 expression is much higher in the bone marrow of acute myeloid leukemia (AML) patients, compared to cancers derived from other tissues (mostly solid tumors), while it has a similar expression in most normal human tissues (Figure 3A). SRRM2 was significantly upregulated in the bone marrow from patients with different AML subtypes compared to healthy samples, highlighting a general SRRM2 overexpression across AML subtypes with diverse underlying genetic alterations (Figure 3B).

**Figure 3.**
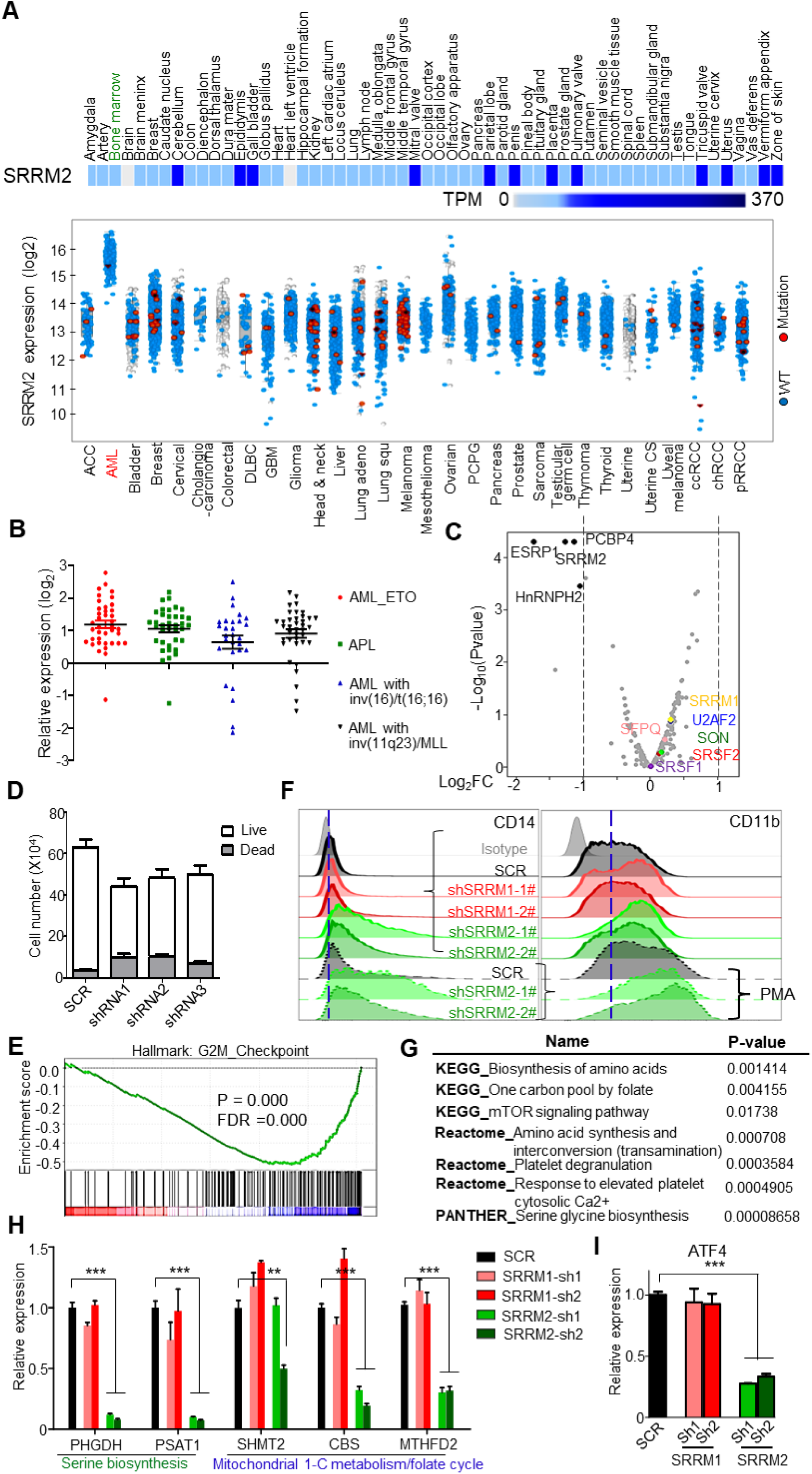
SRRM2 is associated with acute myeloid leukemia. (**A**) Expression of SRRM2 mRNA in human normal tissues (upper heatmap) and cancers (lower graph). The normal tissues display SRRM2 expression from 0 to 370 Transcripts per Million (TPM), to show that SRRM2 was expressed at roughly the same level in most human normal tissues including bone marrow (highlighted in green) (data from The Human Protein Atlas). SRRM2 mRNA levels in cancers (after log2 transformation) show a specific upregulation in AML bone marrow when compared to other tumor or cancers (data from TCGA Research Network: https://www.cancer.gov/tcga). The increased SRRM2 expression in AML is not due to point mutations (very few red dots indicating samples with SRRM2 mutations). (**B**) Expression analyses on patient samples showing SRRM2 upregulation in bone marrow from different types of AML (data from BloodSpot) (57). (**C**) Volcano plot showing log_2_ fold change (log_2_FC) of RNA expression of splicing factors between knockdown and control in THP-1. We marked out with black color the significantly changed (log_2_FC>1, q-value<0.05) splicing factors, and representative splicing factors with other colors. (**D**) Numbers of live/dead cells upon SRRM2 knockdown in THP-1 cells. We cultured 3,000 virus-infected cells (72 h) for 2 days before counting cells. (**E**) GSEA analysis showing that knockdown of SRRM2 influenced G2M checkpoint. We compared gene expression of SRRM2 knockdown to scramble control, based on RNA-seq data. P-value=0.0000, FDR=0.0000. We used hallmark gene sets and gene ontology (GO) gene sets for this analysis. (**F**) FACS assay for detection of the THP-1 surface expression of CD14 and CD11b following shRNA mediated knockdown of splicing factors, without or with PMA treatment (10 ng/ml for 48 h). (**G**) Pathway analysis by KEGG, Reactome, and PANTHER showing that SRRM2 knockdown downregulates one carbon metabolism (serine metabolism) pathways. (**H and I**) Gene expression validation by qRT-PCR showing significant downregulation of genes that govern serine biogenesis (*PHGDH*, *PSAT1*) and mitochondrial 1-C metabolism/folate cycle pathways (*MTHFD2*) (H) and their upstream transcription factor *ATF4* (I), upon SRRM2 but not SRRM1 knockdown. Data are mean ± s.e.m. (n = 3 experiments). Statistical significance determined using one-way ANOVA test (*P < 0.05; **P < 0.01; and ***P < 0.001. ns: no significant difference).

To gain insight into SRRM2 role in myeloid cells and AML, we further performed experiments on the AML cell line THP-1. Immunofluorescence experiments in THP-1 confirmed that nuclear-speckle formation follows the same regulation as in HEK293T (Figure S3). SRRM2 knockdown in THP-1 did not reduce the RNA expression of splicing factors in speckles including *SRRM1*, *U2AF2*, *SRSF1*, *SRSF2*, *SON*, and *SFPQ* (Figure 3C and S4A). The epithelial splicing factor ESRP1 was downregulated upon SRRM2 knockdown, and could account for some of the splicing changes described below. The genes whose expression was affected by SRRM2 knockdown are associated with amino acid and protein metabolism and inflammatory response, indicating that SRRM2 regulates important cellular processes and innate immune function (Figure S4B). We observed a reduction of live cells and an increment of dead cells upon SRRM2 knockdown in THP-1, which was consistent with SRRM2 knockdown influencing the cell cycle and G2M checkpoint (Figure 3D and 3E). This sensitivity to SRRM2 levels appeared specific to naïve THP-1 cells, because we did not see it in either HEK293T cells or PMA-induced differentiated macrophage-like THP-1 cells (data not shown). In addition, SRRM2 knockdown upregulated the surface expression of the mature myelomonocytic markers CD14 and CD11b in both THP-1 and PMA-treated THP-1 cells (Figure 3F and S4C), and also elevated the relative mRNA expression of pan-differentiation markers such as *CD14*, *CD11b*, *CD36*, and *CD163* in PMA induced macrophage-like THP-1 (Figure S4D). Hence, suppression of SRRM2 affects promyelocytic leukemia cell viability and differentiation. One potential mechanism underlying this phenotype is loss of homeostasis on ATF4-serine metabolism pathway, as pathway analysis by KEGG, Reactome, and PANTHER showed that SRRM2 knockdown downregulates one carbon metabolism (serine metabolism) pathways (Figure 3G), including the *PHGDH*, *PSAT1*, and *MTHFD2* genes (Figure 3H, Figure S4E to S4G). Importantly, expression of the master transcription factor for one carbon metabolism *ATF4* was strongly downregulated upon SRRM2 knockdown (Figure 3I), indicating that SRRM2 regulates this pathway via ATF4. Since downregulation of serine metabolism is a crucial metabolic mechanism for AML therapy (35), our results indicate that SRRM2 could be a potential target for the treatment for AML.

### SRRM2 condensates maintain proper alternative splicing in myeloid cells

We carried out transcriptomics upon SRRM2 knockdown in THP-1, as a model/system to link nuclear-speckle disruption with changes in alternative splicing and immune cell function. As a SRRM2 co-regulator (17), we used as control SRRM1, which contains long RS IDRs yet it lacks the NCK and UPR/DPR. We knocked SRRM1/2 down with three different shRNAs, performed RNA-seq and derived the differential alternative splicing events (DASEs) using rMATS (29). Gene ontology showed that the potential splicing targets of SRRM1 or SRRM2 were enriched in regulation of cell cycle progression (Figure 4A). Nevertheless, their respective DASEs were largely different with little overlap, and displayed a different splicing pattern (Figure 4B to 4D). We performed RT-PCR on 47 DASEs of either SRRM1, SRRM2, or both (Figure 4E to 4G), achieving a 66% validation rate using at least two different shRNAs (Figure S5A to S5D), and with splicing change (Δ percentage spliced in or ΔPSI) by RT-PCR highly correlated with that by RNA-seq (Figure 4H, Pearson’s R^2^ = 0.8). To confirm the specificity of SRRM2 targets, we employed exogenous splicing minigenes for either *FES* or *SH3BP2*, and found that minigene splicing was disrupted upon SRRM2 knockdown much like the corresponding endogenous DASEs (Figure 4I). These results indicate that joint SRRM1/2 regulation is not the major mechanism of SRRM2-mediated alternative splicing.

**Figure 4.**
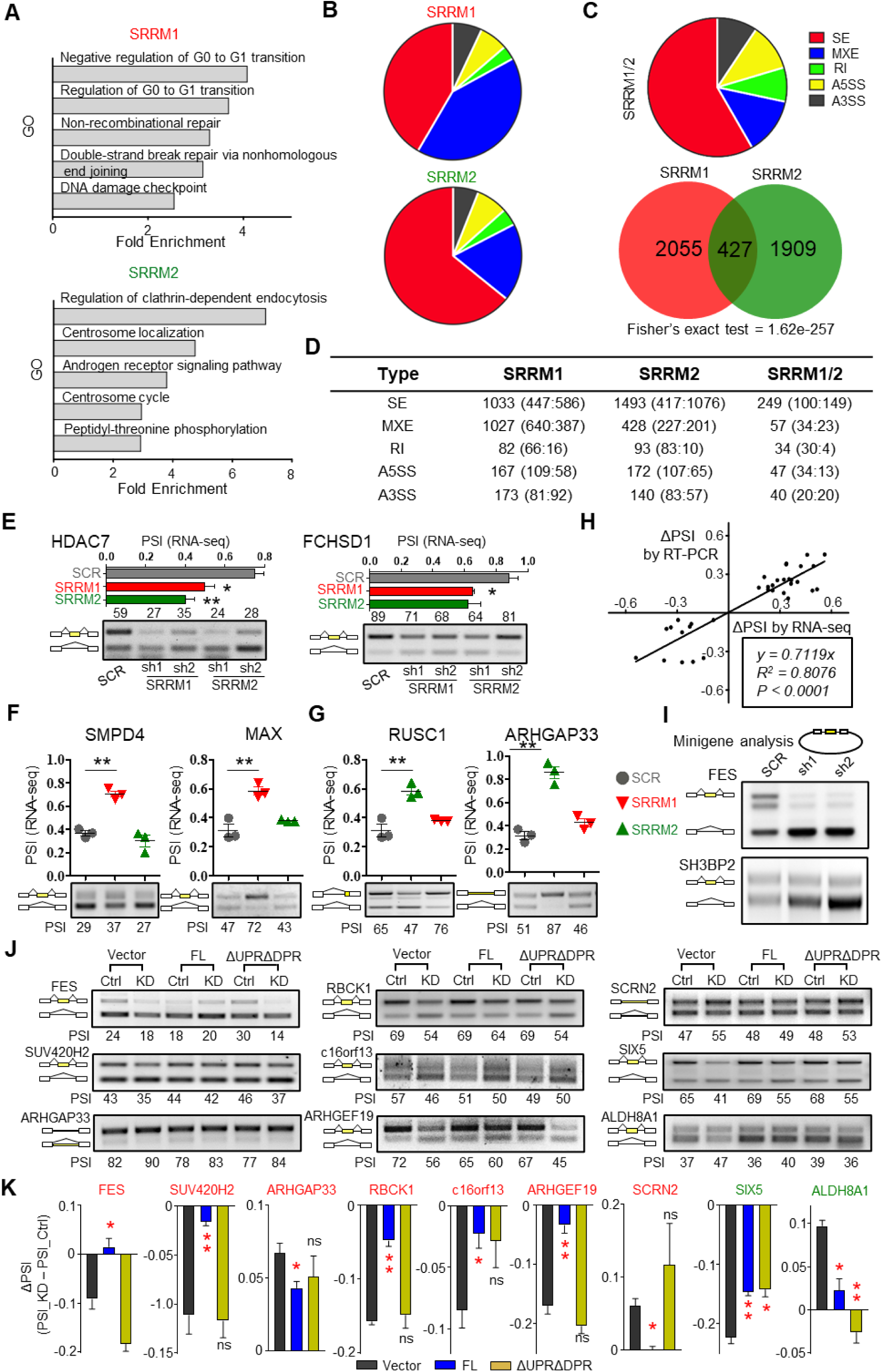
SRRM2 condensates maintain proper alternative splicing. (**A**) Gene Ontology of DASEs induced by SRRM1 or SRRM2 knockdown. (**B**) Splicing subtypes of DASEs induced by SRRM1 or SRRM2 knockdown. (**C**) Overlap and subtypes of SRRM1 and SRRM2 co-regulated DASEs. SE: skipped (cassette) exon; MXE: mutually exclusive exon; RI: Retained Intron; A5SS: alternative 5’ splice sites; A3SS: alternative 3’ splice sites. (**D**) Numbers of SRRM1 and SRRM2 regulated DASEs. We show for each splicing subtype the numbers of total, positive ΔPSI (enhanced inclusion or longer isoform) and negative ΔPSI (enhanced skipping or shorter isoform). (**E**) Two validation examples of SRRM1/SRRM2 co-regulated DASEs. Upper graph: PSI based on rMATS analysis. Lower gel: RT-PCR validation showing percentage of inclusion band. (**F**) Validation examples of SRRM1 (red) but not SRRM2 (green) DASEs. (**G**) Validation examples of SRRM2 (green) but not SRRM1 (red) DASEs. (**H**) Correlation of ΔPSI between RT-PCR validation and rMATS analysis of RNA-seq data. (**I**) Minigene experiment of SRRM2 targets *FES* and *SH3BP2*. We transfected either control or SRRM2 knockdown HEK293T cells with minigenes and visualized minigene splicing patterns via RT-PCR. DASEs (|ΔPSI|>0.1, FDR<0.05). (**J and K**) DASE analysis by RT-PCR (J) and quantification of ΔPSI (PSI_KD-PSI_Ctrl) (K) upon SRRM2 knockdown and rescue with Control, Full-length (FL) SRRM2, or ΔUPRΔDPR. Red label: genes with splicing change caused by KD can only be rescued by FL; green label: targets rescued by both FL and ΔUPRΔDPR. Data are mean ± s.e.m. (n = 3 experiments). Statistical significance determined using one-way ANOVA test (*P < 0.05; **P < 0.01; and ***P < 0.001. ns: no significant difference).

Importantly, the UPR/DPR double deletion disrupted SRRM2 condensates (Figure 2B), yet these repeat regions were essential for most but not all SRRM2 splicing targets: the splicing of *SIX5* and *ALDH8A1* was rescued by both overexpressed wild-type or ΔUPRΔDPR mutant, while the splicing of *FES* and six other targets was only rescued by the wild-type SRRM2 (Figure 4J and 4K). These results indicate that alternative splicing dysregulation upon SRRM2 knockdown is partly but not entirely associated with its condensate formation.

### Features of SRRM2 splicing targets

SRRM2 knockdown induced a significantly increased number of exon-skipping events when compared to SRRM1 (Figure 5A). Splice-site strength estimation (36) revealed that SRRM2 knockdown tends to induce skipping of cassette exons with weak 5’ splice sites, followed by downstream exons with strong 3’ splice sites (Figure 5B). The introns both upstream and downstream of cassette exons skipped by SRRM2 knockdown were on average 5.5 kb shorter than the introns with increased inclusion upon knockdown, and control introns as well (Figure 5C). Motif analysis using MEME (37) revealed that the cassette exons skipped by SRRM2-knockdown are enriched with the GGUGG motif either in the intronic or exonic regions (Figure S6A). Overall, these analyses reveal the specific features exhibited by SRRM2-regulated DASEs (Figure 5D).

**Figure 5.**
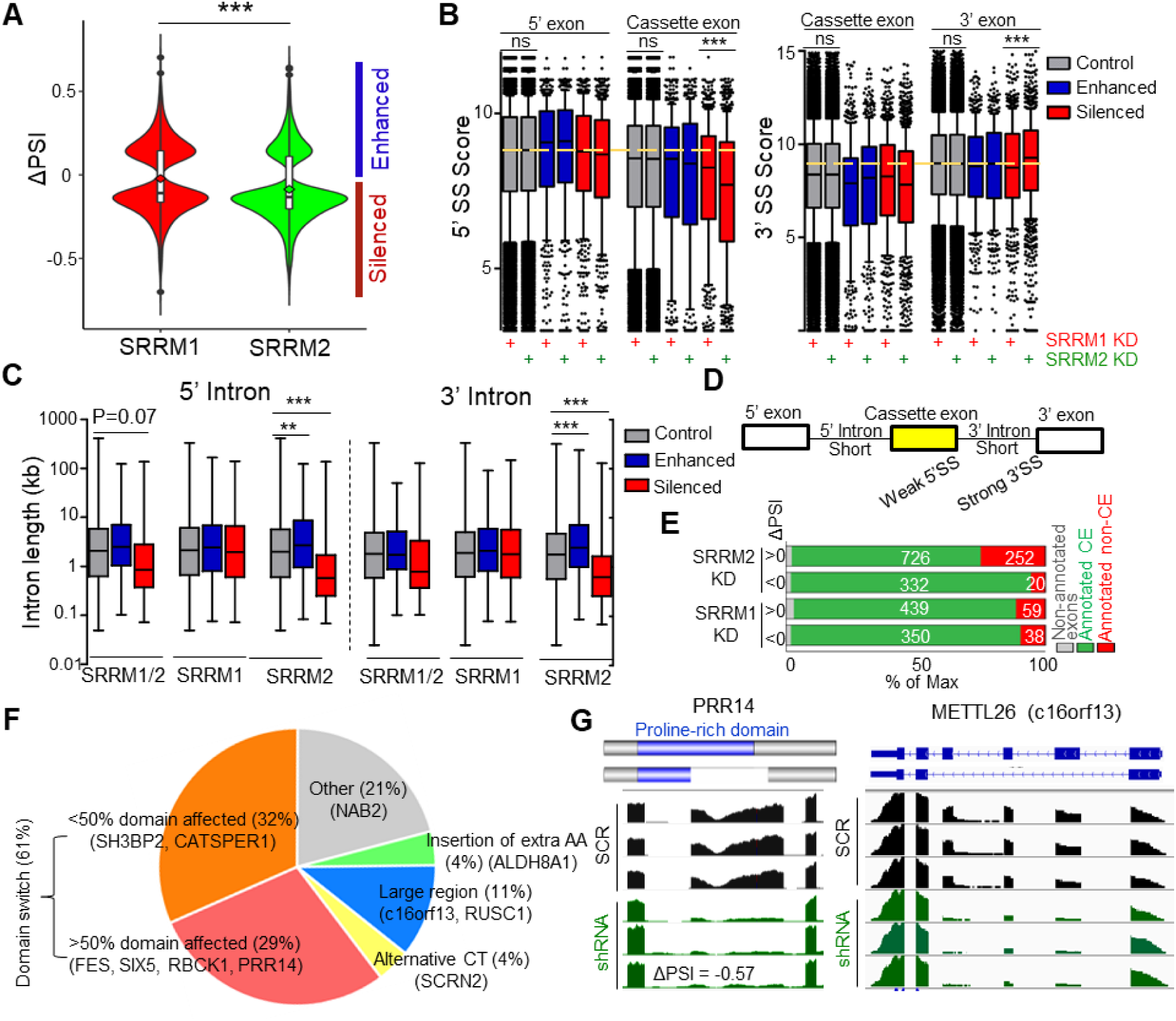
SRRM2 targets exhibit specific splicing features. (**A**) Violin plots showing that knockdown of SRRM2 mainly induced exon skipping, with SRRM1 knockdown as control. Enhanced/Silenced: higher inclusion/skipping of cassette exons. (**B**) Splice-site strength of SRRM2-regulated DASEs. We analyzed the 5’ SS (splice site) score of 5’ exons and cassette exons and 3’ SS score of cassette exons and 3’ exons of either SRRM1- or SRRM2- regulated DASEs. Yellow lines label the average score of controls. (**C**) Intron length of SRRM2-regulated DASEs, compared to either SRRM1 or SRRM1/SRRM2 coregulated DASEs. (**D**) Summary of splicing features of SRRM2. Top, SRRM2 regulated cassette exons possess weak 5’ splice sites and are flanked by short introns. (**E**) SRRM2 knockdown induced exon-skipping events have a higher percentage (25%) of non-annotated cassette exons (CE). (**F**) Domain analysis for validated DASEs upon suppression of SRRM2, with some genes as examples. 61% of validated targets displayed domain change (32% with <50% domain change affected plus 29% with >50% domain change), and 40% have strong protein changes (29% of >50% domain change plus the 11% for large region). Alternative CT: alternative C-terminus; AA: amino acid; Large regions: several domains affected concurrently. (**G**) Examples of protein domain change by single-exon skipping of *PRR14* exon 7, and large region removal by multiple skipping of *METTL26* (c16orf13) exons 2 to 5. We show the reads based on RNA-seq data as generated by IGV software. Validation shown in Figure S5A (with PSI derived from RNA-seq and alternative splicing analysis). Statistical significance determined using one-way ANOVA test (*P < 0.05; **P < 0.01; and ***P < 0.001. ns: no significant difference).

Inclusion of cassette exons with weak splice sites is usually influenced by transcriptional rate (38). To test the effects of SRRM2 in co-transcriptional splicing, we treated THP-1 cells with two drugs that slow down Pol II transcription, 5,6-Dichloro-1-β-D-ribofuranosylbenzimidazole (DRB) and camptothecin (CPT) (38, 39). RNA-seq global assay showed that SRRM2 knockdown enhanced the effect of CPT/DRB induced alternative splicing change on SRRM2 targets (Figure S6B), indicating that SRRM2 tends to affect splicing in the same direction as processive Pol II. Interestingly, nearly 25% of the skipped events upon SRRM2 knockdown were unannotated (Figure 5E and S6C). More than 40% of validated DASEs upon SRRM2 suppression were associated with strong protein change (29% with >50% domain affected, plus 11% with large region removed by splicing) (Figure 5F). As examples, we selected skipping of *PRR14* exon 7 for protein domain change, and skipping of *METTL26* (c16orf13) exons 2 to 5 for removal of large region by splicing (Figure 5G and S5A). Domain changes, and especially large region removal by multiple exon skipping is a significant mechanism for protein dysfunction (40). Thus, alternative splicing is the major mechanism for the maintenance of protein function and cell homeostasis by SRRM2.

### Function of SRRM2 splicing targets in myeloid cells

Next, we investigated the functional impact of the SRRM2-regulated alternative splicing. The FES SH2 domain encoded by cassette exon 10 is essential for attenuating the TLR signaling pathway (Figure S7A) (41, 42), and this exon was significantly excluded upon SRRM2 knockdown in either monocytic or macrophage-like THP-1 cells (Figure 6A and S5D). We further tested the isoform-specific function of FES using shRNA targeting either long isoforms only (with exon 10) or both short and long isoforms (Figure 6B and 6C) on PMA-induced macrophage-like THP-1. We found that the *TNFα* and *IL1β* pro-inflammatory cytokines were significantly upregulated upon specific knockdown of all isoforms and SH2-containing *FES* isoforms (Figure 6D). The shRNA targeting all isoforms more strongly upregulated these cytokines, possibly due to its higher knockdown efficiency than the exon-10 specific shRNA (Figure 6C), which could not be further optimized due to the limited length of exon 10. Hence, the exon-10 containing *FES* isoforms contribute to the observed transcriptional modulation of proinflammatory cytokines in macrophages by SRRM2 (Figure S7B). This may partially explain why human immune cells are more sensitive to LPS, as mice only have the FES isoform with SH2 domain, while human macrophages exhibit significant skipping of exon 10 (Figure 6B). Collectively, these data suggest that SRRM2 attenuates innate inflammatory responses via *FES*, although there might be other factors and pathways involved.

**Figure 6.**
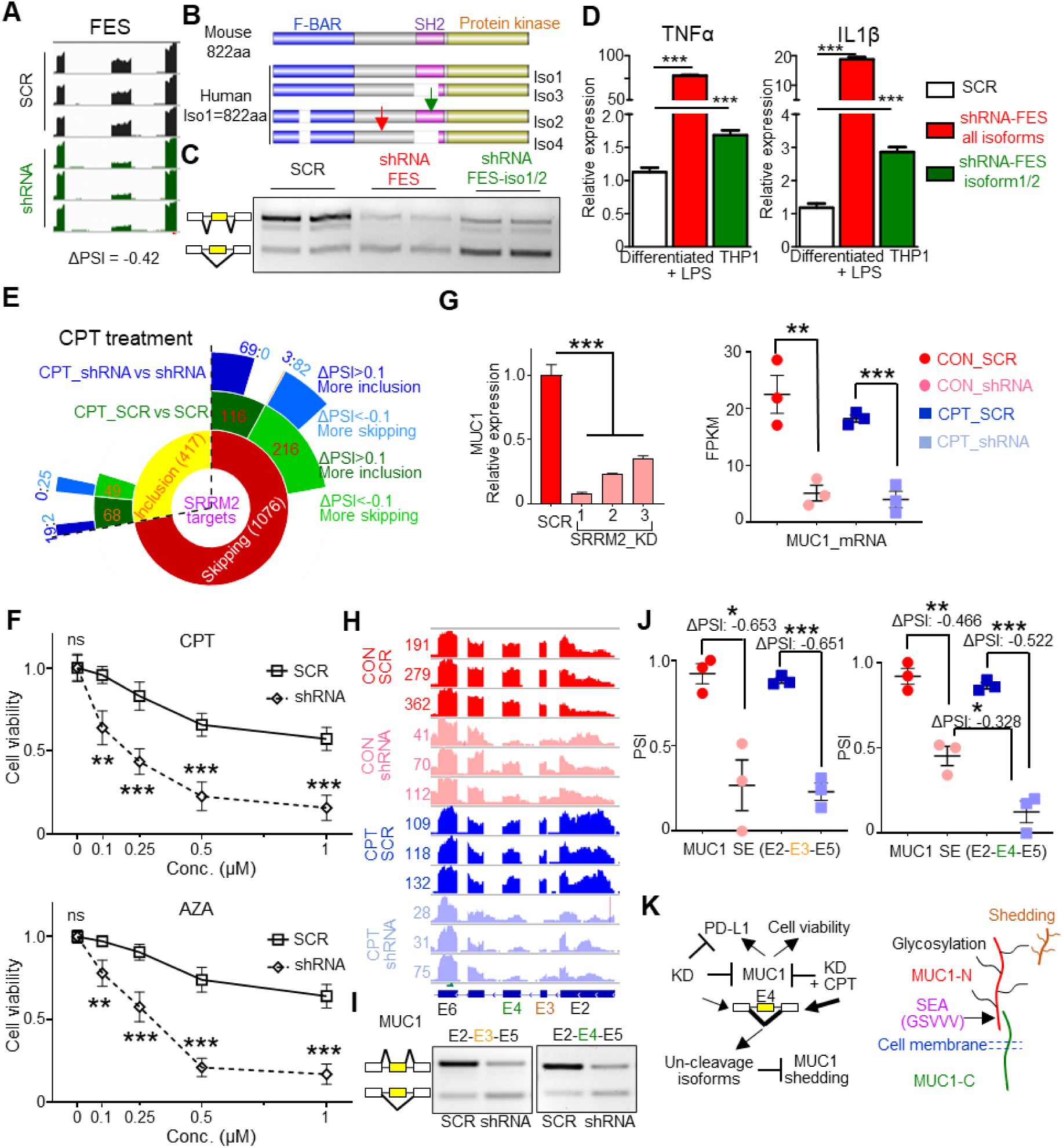
Alternative splicing of the SRRM2 targets *FES* and *MUC1* regulates innate immunity and cell homeostasis. (**A**) SRRM2 regulates alternative splicing of *FES*. Map of reads: IGV software; PSI: rMATS analysis; Splicing validation by RT-PCR (Figure S5A). (**B**) Diagram of mouse and human FES domains including SH2 affected by alternative splicing in human only, and targeting shRNAs (green arrow showing shRNA targeting iso1/2, red arrow showing shRNA targeting all isoforms). Iso, isoform. (**C**) Isoform-specific knockdown effect of FES shRNAs. (**D**) Relative expression of *TNFα* and *IL1β* proinflammatory cytokines upon isoform-specific knockdown of *FES* in human macrophage-like THP-1 cells under LPS (1 ng/ml) for 6 h. To observe a stronger phenotype upon *FES* knockdown, we used a weak LPS stimulation with little effect on THP-1 inflammation. (**E**) Sunburst diagram showing the effect of CPT drug, and combined CPT plus SRRM2 knockdown on SRRM2 DASEs. (**F**) Cell viability analysis of THP-1 cells upon SRRM2 knockdown with either CPT or AZA treatment for 24 h. (**G**) Expression of MUC1 upon SRRM2 knockdown in THP-1, measured by RNA-seq or qRT-PCR. (**H**) RNA-seq analysis showing the reads and annotations of spliced exons of *MUC1* upon SRRM2 knockdown and CPT treatment. (**I**) RT-PCR validation of the skipping of exons 3 or 4 upon SRRM2 knockdown. (**J**) PSI differences and statistical analysis of *MUC1* exon 3 and 4 upon SRRM2 knockdown and combined treatment with CPT. (**K**) Graphic diagram showing SRRM2 regulates exon 4 skipping to modulate MUC1 shedding. MUC-N: N-terminal domain; MUC1-C: C-terminal domain; SEA: Sperm protein, Enterokinase and Agrin; GSVVV: cleavage site for proteases TACE or ADAM9; Glycosylation: protein modification. Shedding generates soluble MUC1, promoting cancer progression or regulating immune response. Data are mean ± s.e.m. (n = 3 experiments). Statistical significance determined using one-way ANOVA or two-way ANOVA (F) with Bonferroni correction for multiple comparison. (*P < 0.05; **P < 0.01; and ***P < 0.001. ns: no significant difference).

We showed that CPT drug treatment as a transcription inhibitor influences alternative splicing in THP-1 cells. Importantly, CPT and its derivatives are drugs for various cancers including leukemia (43). Consistently, GO analysis shows that CPT treatment slows down Pol II and induces cell cycle arrest and programmed cell death (Figure S7C). SRRM2 knockdown enhanced the CPT drug treatment effect on SRRM2 DASEs (Figure 6E). Consistent with this, when we combined SRRM2 knockdown with either CPT or azacytidine (AZA) treatment, we found that THP-1 cell viability was significantly lower upon combination treatment compared to drug alone (Figure 6F). The DEGs or DASEs of CPT/SRRM2 knockdown compared to CPT alone were enriched in the pathways of cell cycle arrest and DNA damage (Figure S7D). Among the DASE gene list, the oncoprotein MUC1 (Mucin-1) is a potential AML therapeutic target (44). Here we show that SRRM2 knockdown significantly downregulated MUC1 at both mRNA and cell surface expression level (Figure 6G, S7E, and S7F). SRRM2 knockdown significantly induced strong skipping of either exon 3 or exon 4 in THP-1 cells (Figure 6H and 6I), with the combination treatment inducing more skipping of exon 4 compared to either SRRM2 knockdown or CPT alone (Figure 6J). Disruption of the GSVVV cleavage site in MUC1 SEA (Sperm protein, Enterokinase and Agrin) domain encoded by exon 4 prevents shedding of MUC1 mediated by proteases such as TACE or ADAM9, and MUC1 shedding is a mechanism to promote cancer progression, as well as it regulates innate immune response (45, 46) (Figure 6K and S7F). Thus, the targeting MUC1 by SRRM2 contributes to the reduction of cell viability by AML drugs.

## DISCUSSION

This study provides evidence for SRRM2 as the main organizer or scaffold of nuclear-speckle formation, and that this is important for alternative splicing and innate immunity. While knockdown of many components changes nuclear-speckle number, shape, and composition, SRRM2 is in the core and essential for organizing speckles. Disease-associated tau aggregates disrupt the spatial organization of speckles, coacervating SRRM2 (but not SON) into the center of the aggregates, and accumulating mainly SRRM2 (but not SON or SRSF2) in the cytosol (34). Hence, splicing-speckle proteins should be clients that partition into speckles via SRRM2. Here we show that SRRM2 condensates exhibit the following phase separation hallmarks (8): spherical shape, dynamic rearrangement and coalescence, both in cells under controlled expression by knock-ins and with two small fragments *in vitro*, as well as concentration dependence for the two *in vitro* fragments. In addition to the lack of *in vitro* data for full-length SRRM2 because of its enormous size, we are only missing the diffusion across boundary criterion, whose information helps distinguish phase separation over other phenomena such as polymer scaffold binding or bridging. Nevertheless, our work supports the notion that SRRM2 is to nuclear speckles the main organizational or scaffold, as a counterpart of G3BP to stress granules (47), or nucleophosmin to nucleoli (48). SRRM2 organizes condensates very likely via phase separation.

We showed that SRRM2 has an intrinsic propensity to form condensates via multiple IDRs, whose distinct physicochemical properties we partially characterized. Phase Separation driving forces include cation-π interactions between tyrosine and arginine, and coulombic interactions between repeated positively-charged lysine or arginine with negatively-charged nucleic acids (49, 50). We found that KSR and three 11-amino acid repeats of UPR are sufficient to form condensates *in vitro* with different properties, while UPR/DPR are essential for speckle formation. Based on these results and the highly repetitive nature of SRRM2 IDRs, most small deletions and point mutations in these domains are not expected to change the condensation (and splicing) properties of SRRM2. Moreover, as arginine-rich mixed charge domains target proteins to speckles while lysines target to nucleolus (50), we postulate that the extensive SRRM2 RS domains and other IDRs dominate over the lysines in KSR. The SRRM2-IDR coacervation with RNA and droplet formation propensity under mimicked intracellular molecular crowding, suggest that SRRM2 interplays with splicing-speckle proteins such as the scaffold SON, and with unspecific pre-mRNA or non-coding speckle RNA MALAT1. The future characterization of the condensate properties in cells and *in vitro* for all SRRM2 IDRs, individually and in combinations, should shed light on the nature and dynamics of SRRM2 condensates, and link them to alternative splicing. Encouragingly, we already found that UPR/DPR are differentially linked to regulation of splicing targets, as a functional connection between alternative splicing and condensate formation. We speculate that the conserved NCK might help dissolve ΔUPRΔDPR to localize to the spliceosome, which is enough for some but not all SRRM2 targets. Further experiments will elucidate the IDR and condensate dependence of SRRM2-regulated splicing events.

Live-cell tracking during mitosis strongly supported a major role of SRRM2 in speckle formation. SRSF2 enters the newly formed nucleus and stays in a diffuse pattern during cytokinesis, while SRRM2 enters later concomitant to speckle formation. In cell cycle, speckle dissolution, MIG formation, and speckle re-nucleation depend on post-translational modifications. Previous studies demonstrated that DYRK3, but not other kinases such as SRPK1 or CDK1, interacts with SRRM2 and controls MIG formation (7). In early phase Alzheimer’s disease brains, ERK prevented SRRM2’s nuclear translocation by inhibiting its interaction with TCP1α (T-complex protein subunit α), disrupting PQBP1’s localization and targets (20). Modulation of SRRM2 dynamics by kinases under physiological condition remains unclear, yet candidates are cyclin-dependent kinases (CDKs) or CLK family members. Possible phosphorylation sites are the RS dipeptides in the RS, KSR or UPR/DPR. Both phosphorylated-tau and SRRM2 exhibit phase separation properties, and the positively-charged IDRs in SRRM2 characteristically form aggregates when there is no cell-cycle induced phosphorylation modification on cytosolic SRRM2 in terminal differentiated neurons (34, 51). Pol II Carboxy-Terminal Domain (CTD) with heptad-repeats exhibits specific phosphorylation patterns in promoter-associated transcription initiation (hypo-phosphorylation) and elongation (serine-2 hyperphosphorylation) by CDKs, to form distinct condensates (52, 53). As parallel to CTD heptad repeats, the R/S-containing UPR/DPR repeats may play key roles in SRRM2 condensate properties and dynamics, consistent with the essentiality of both IDRs for nuclear-speckle formation, and with the SRRM2’s featured role in alternative splicing.

Few studies are linking phase separation with immune functions: for instance, *in vitro* reconstitution of membrane complexes to recapitulate T-cell activation established that phase separation is crucial for the kinetic proofreading regulation of Son of Sevenless (SOS), by a slow recruitment and enzymatic activation (54). Our results suggest that splicing condensates maintain innate immune-cell homeostasis in multiple aspects including one carbon metabolism/folate cycle and serine metabolism, cell viability, differentiation, and innate immune response. Hence, the link between nuclear speckles and immune homeostasis is to some extent dependent on alternative splicing and SRRM2 via its condensation properties.

Mutations of splicing factors are linked to a wide range of solid cancers and leukemia (55). We show that deficiency of SRRM2 leads to a defect in cell viability along with skipping of cassette exons with specific features, and abnormal expression of SRRM2 without mutation is found in AML patients, suggesting that its expression level is important for maintaining proper cell homeostasis including alternative splicing in AML cells. Furthermore, the SRRM2-mediated regulation of cassette exon inclusion, by a mechanism independent of RRM, could be achieved by the biophysical properties of SRRM2 condensates, likely via phase separation. This is consistent with the regulatory mechanism of other splicing factors like Rbfox2, which serves as a scaffold to recruit other components, so as to form aggregates to regulate exon skipping (10). The localization of SRRM2 and other splicing factors may contribute to splicing decisions, as in the model proposing that sequence-dependent RNA positioning along the nuclear speckle interface coordinates RNA splicing (56). Further studies on the regulatory mechanism of SRRM2 in alternative splicing may connect the biophysical properties of the condensates with their function in splicing.

SRRM2-induced exon skipping caused a high percentage of protein domain change and removal of large regions. The extent to which the concomitant ESRP1 reduction contributes to the SRRM2 targets will be determined in the future. We characterized the alternative splicing of *FES* as an example for innate immune response, and the AML therapeutic target *MUC1*. Other SRRM2-regulated targets with functional relevance are *PRR14*, whose domain change might affect tumor cell proliferation, and *NAB2*, whose change might help overcome the differentiation block in leukemia. Further studies should elucidate the SRRM2 targets in relation to other key regulators of AML and myeloid cell homeostasis.

## DATA AVAILABILITY / ACCESSION NUMBERS

All sequencing data is available in the Gene Expression Omnibus (GEO) under the series accession number GSE199720. Flow cytometry data is available in FlowRepository with the accession number FR-FCM-Z56E.

## SUPPLEMENTARY DATA

Supplementary Data are available at NAR Online.

## ACKNOWLEDGEMENT

We thank Carl Zeiss for providing the ideal platform for confocal 4D imaging (LSM 980 with Airyscan 2). Oliver Mueller-Cajar provided helpful comments on the manuscript.

## Author contributions

XR supervised the project. SX and XR conceived the study and designed the experiments. SX performed most experiments and in silico analyses with help from S-KL (microscopy), DYS (AS validation and some in silico analyses), WSLA (protein purification). HYL provided suggestions. SX and XR wrote the paper.

---

**Graphical abstract.**
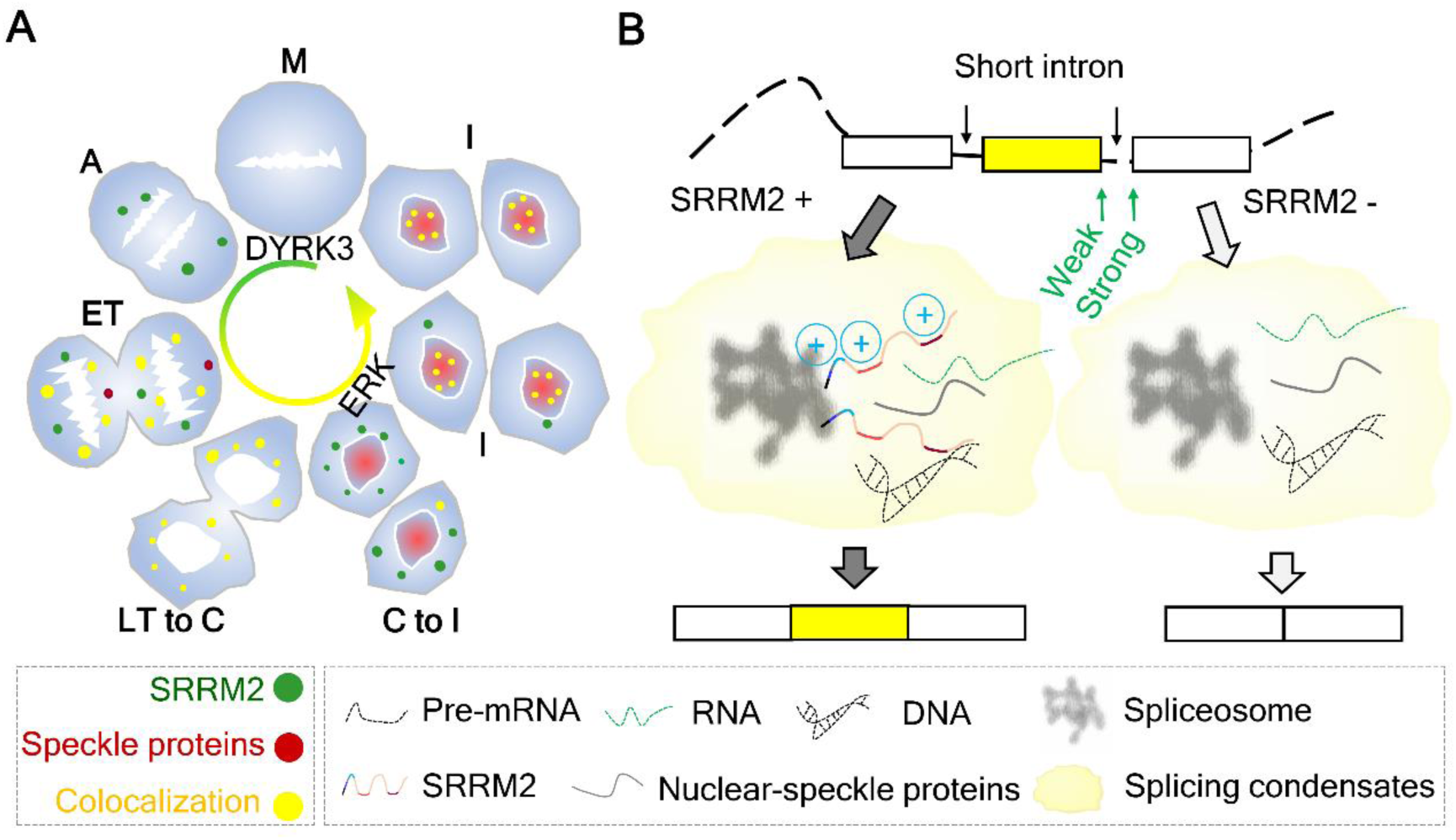
Schematics of splicing-condensate organization and alternative splicing regulation by SRRM2. (A) Dynamics of nuclear-speckle proteins during cell cycle. M: Metaphase (low DYRK3), SRRM2 and SRSF2 are dispersed throughout cells. A: Anaphase, SRRM2 starts to form condensates. ET: Early Telophase, SRSF2 starts to form condensates, mostly colocalizing with SRRM2 condensates. LT to C: Late Telophase to Cytokinesis, stage 1: nearly all speckle proteins colocalize in MIGs; stage 2: SRSF2 dissolves from MIGs and binds to chromatin. C to I: Cytokinesis to Interphase, stage 1: condensed SRRM2 is dissolved and imported into the new nucleus, to form nuclear-speckle condensates; stage 2: intensity and size of nuclear-speckle condensates increase with SRRM2 intensity. (B): Splicing condensates maintain proper alternative splicing and innate immune homeostasis. Three SRRM2 IDRs promote condensates to induce exon inclusion, while SRRM2 deficiency induces skipping of cassette exons with short introns and weak 5’ splice sites.

