## Supplementary data for "SRRM2 organizes splicing condensates to regulate alternative splicing"

#### Supplementary Figures

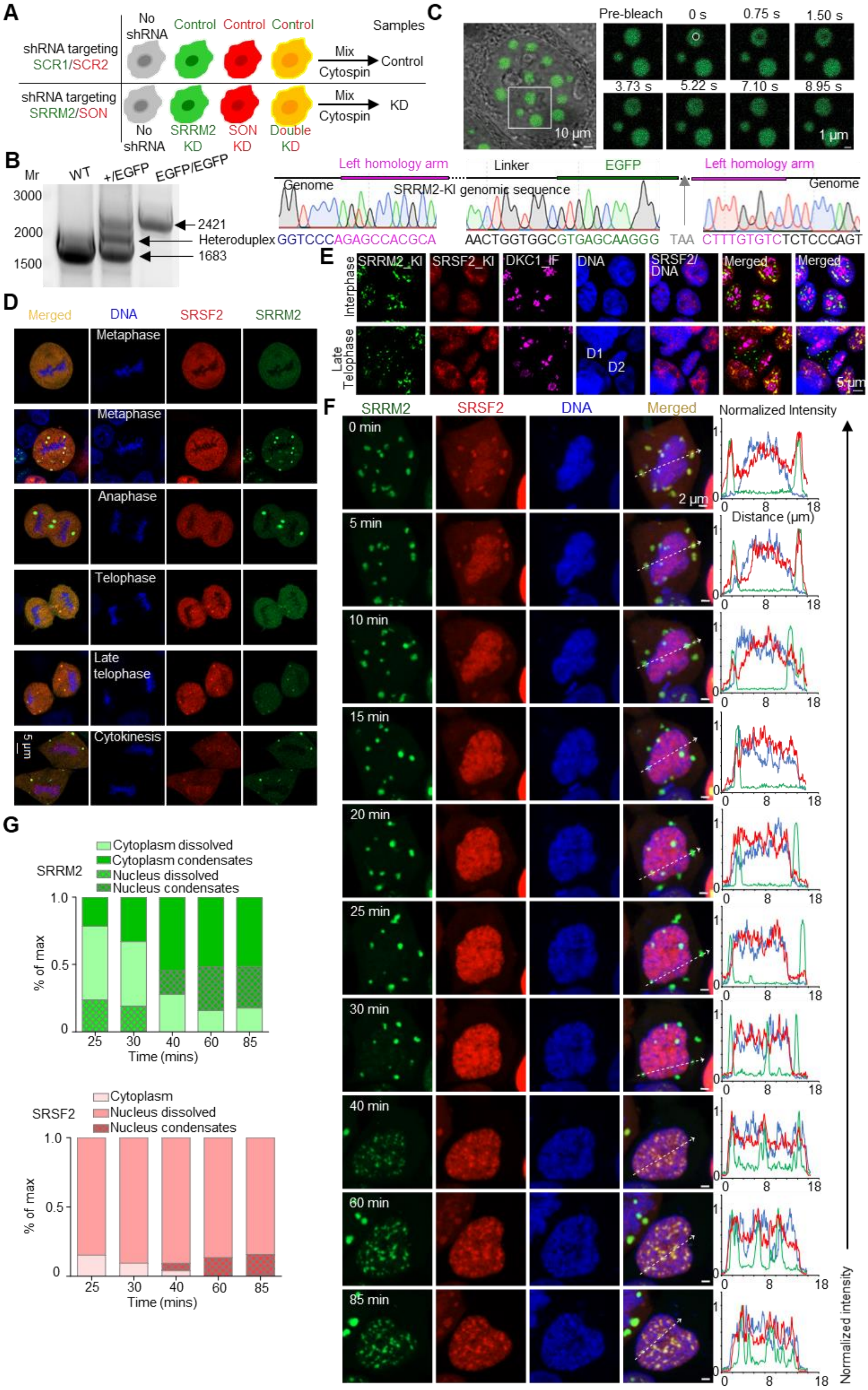

**Figure S1. SRRM2 is responsible for organizing nuclear speckles**

(**A**) Schematic of the experimental design for the shRNA mediated double knockdown (KD) of SRRM2 and SON in HEK293T cells. (**B**) Validation of knock-in cells by RT-PCR genotyping and DNA sequencing including entire recombination arms and knock-in regions (SRRM2 as an example). (**C**) Representative images of FRAP experiments on overexpressed EGFP-SRRM2 in HEK293T cells. (**D**) Images of HEK293T cells co-transfected with EGFP-SRRM2 and mCherry-SRSF2 showing distribution of SRRM2 and SRSF2 in different cell cycle stages. (**E**) Immunofluorescence staining on fixed double knock-in cells showing the distribution of DKC1 in late telophase and interphase. (**F**) Representative 4D (real-time 3D) live-cell imaging (maximum intensity projection of stacks) for tracking SRRM2 and SRSF2 localization from cytokinesis to interphase. See Video 2. Intensity: the normalized fluorescence intensity of condensed/dissolved SRRM2 (green)/SRSF2 (red) and Hoechst (blue) for the line scan analysis. (**G**) Percentage of dissolved/condensed SRRM2 or SRSF2 in cytoplasm or nucleus after formation of new nucleus.

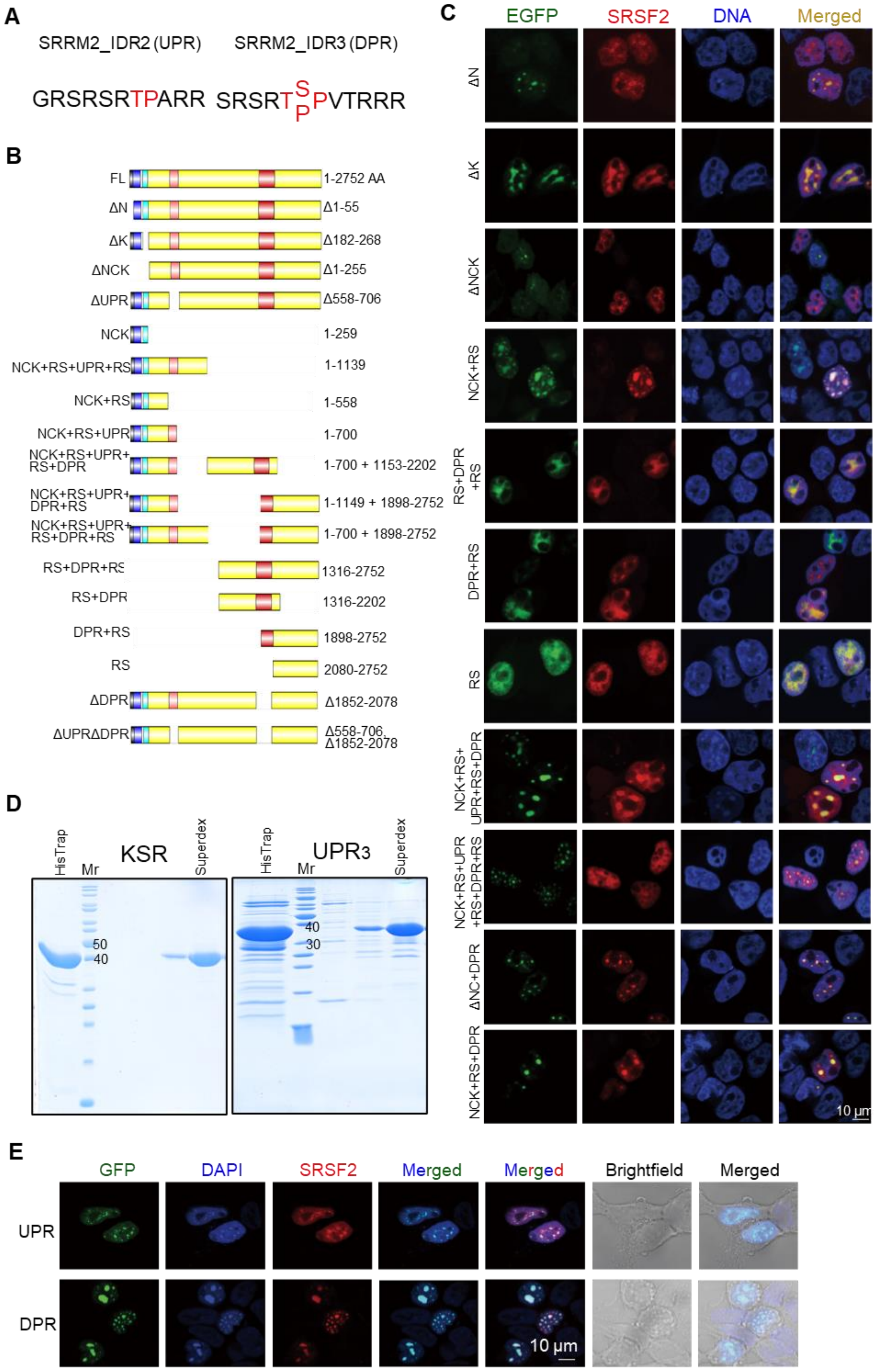
A
SRRM2_IDR2 (UPR)
SRRM2_IDR3 (DPR)
GRSRSRTPARR
SRSRT S P PVTRRR
B
FL
1-2752 AA
ΔN
Δ1-55
ΔK
Δ182-268
ΔNCK
Δ1-255
ΔUPR
Δ558-706
NCK
1-259
NCK+RS+UPR+RS
1-1139
NCK+RS
1-558
NCK+RS+UPR
1-700
NCK+RS+UPR+RS+DPR
1-700 + 1153-2202
NCK+RS+UPR+DPR+RS
1-1149 + 1898-2752
NCK+RS+UPR+RS+DPR+RS
1-700 + 1898-2752
RS+DPR+RS
1316-2752
RS+DPR
1316-2202
DPR+RS
1898-2752
RS
2080-2752
ΔDPR
Δ1852-2078
ΔUPRΔDPR
Δ558-706, Δ1852-2078
C
EGFP
SRSF2
DNA
Merged
ΔN
ΔK
ΔNCK
NCK+RS
RS+DPR +RS
DPR+RS
RS
NCK+RS+ UPR+RS+DPR
NCK+RS+UPR +RS+DPR+RS
ΔNC+DPR
NCK+RS+DPR
10 μm
D
HisTrap
Mr
KSR
Superdex
50
40
HisTrap
Mr
UPR3
Superdex
40
30
E
GFP
DAPI
SRSF2
Merged
Merged
Brightfield
Merged
UPR
DPR
10 μm

**Figure S2. Domain mapping and properties of SRRM2 IDRs**

(**A**) UPR and DPR consist of 12 repeats each of the indicated sequence. (**B**) Diagram of domain mapping of human SRRM2 mutants fused to EGFP. (**C**) Colocalization images of the SRRM2 mutants with SRSF2. We co-transfected HEK293T cells with EGFP-tagged SRRM2 mutants and mCherry-SRSF2. We fixed cells and labeled DNA with DAPI. (**D**) Coomassie blue staining of SDS-PAGE, showing the purity of KSR and $UPR_3$ fused to EGFP. We purified these fusion proteins with HisTrap column followed by gel filtration using Superdex column. (**E**) Images of HEK293T cells co-transfected with either EGFP-UPR or EGFP-DPR fragments and mCherry-SRSF2. DAPI stained for DNA or high concentration of RNA. Brightfield images of transfected cells showing that DPR tended to accumulate and form gel-like condensates or aggregates inside nucleus.

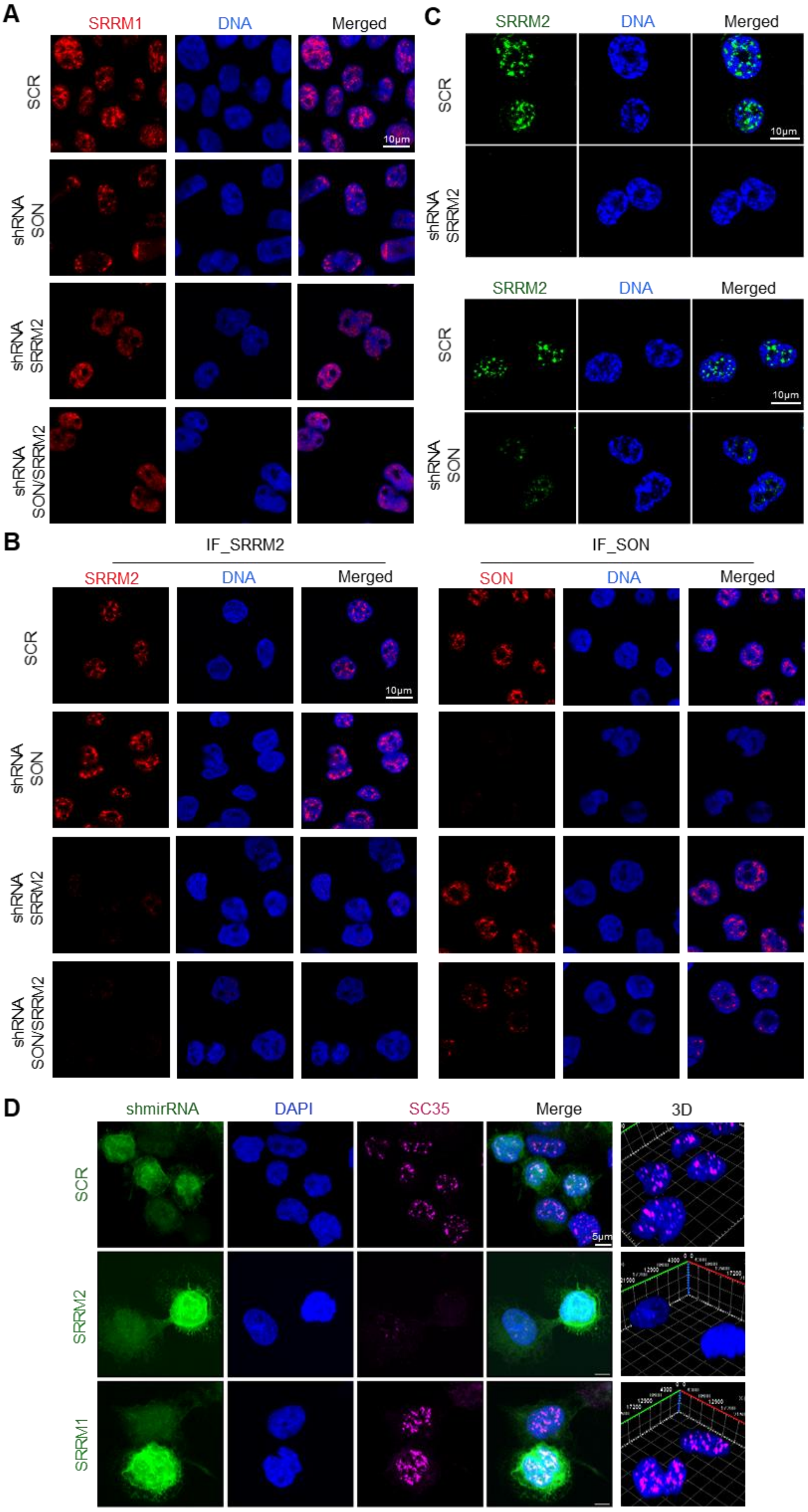

A
SRRM1
DNA
Merged
SCR
shRNA SON
shRNA SRRM2
shRNA SON/SRRM2
10μm
C
SRRM2
DNA
Merged
SCR
shRNA SRRM2
SRRM2
DNA
Merged
SCR
shRNA SON
10μm
B
IF_SRRM2
SRRM2
DNA
Merged
IF_SON
SON
DNA
Merged
SCR
shRNA SON
shRNA SRRM2
shRNA SON/SRRM2
10μm
D
shmirRNA
DAPI
SC35
Merge
3D
SCR
SRRM2
SRRM1
5μm

**Figure S3. Nuclear-speckle staining in THP-1 and differentiated macrophage-like THP-1 cells**

(**A and B**) Representative images of SRRM1 (A), SRRM2 (left panel in B), or SON (right panel in B) immunofluorescence staining upon single knockdown of SRRM2, SON, or SRRM2/SON double knockdown in THP-1 cells. (**C**) Representative images of SRRM2 (top) or SON (bottom) immunofluorescence staining in human macrophage-like THP-1 cells, with shRNA mediated knockdown to show the antibody specificity in the cells. (**D**) Immunofluorescence (IF) showing knockdown of SRRM2 by shmirRNA disrupts nuclear-speckle maker SRSF2 but not SRRM1 in human macrophage-like THP-1 cells, indicating that the SRSF2 antibody (SC35) mainly stains for SRRM2. Co-expressed TurboGFP visually marked knockdown cells. We used both scramble and SRRM1 shmirRNA as controls. Right column, processed images from 3D imaging.

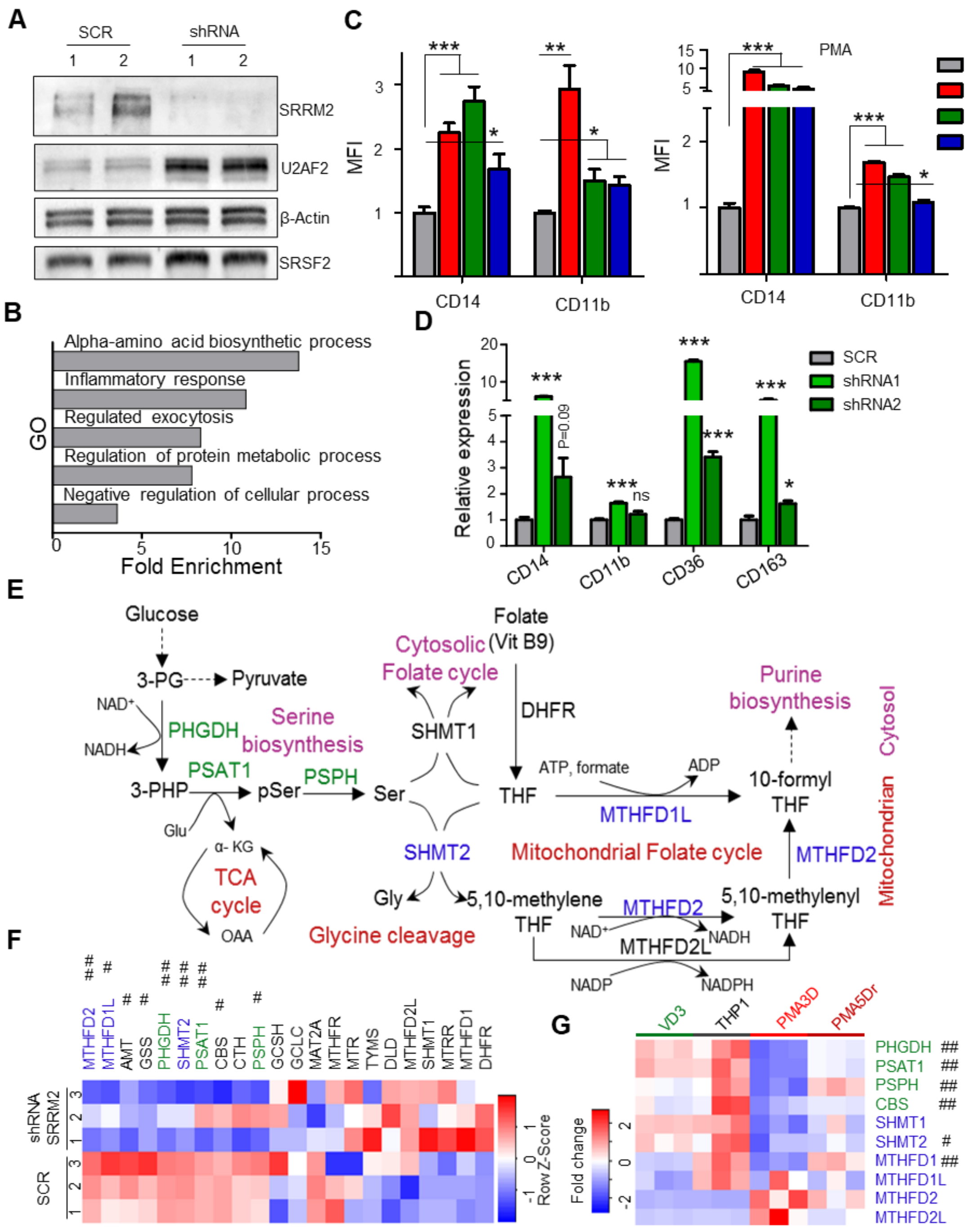

**Figure S4. SRRM2 is a potential target for the treatment of acute myeloid leukemia**

(**A**) Western blot showing protein expression of SRRM2, U2AF2, and SRSF2 upon SRRM2 knockdown in THP-1. (**B**) GO analysis showing SRRM2 knockdown in THP-1 induced differently expressed genes (DEGs). (**C**) Relative expression level of CD14 and CD11b on THP-1 upon knockdown of SRRM2 with 3 different shRNAs. MFI: mean fluorescence intensity. (**D**) qRT-PCR showing the relative expression levels of marker genes *CD14*, *CD11b*, *CD36*, and *CD163* in THP-1 upon knockdown of SRRM2. (**E**) Graphic figure showing the regulation of serine metabolism and mitochondrial 1-C metabolism/folate cycle. (**F**) Heatmap showing the expression of genes in serine metabolism and mitochondrial 1-C metabolism/folate cycle upon SRRM2 knockdown in THP-1 cells. (**G**) Heatmap showing the expression of key genes in one carbon metabolism pathways (*PHGDH*, *PSAT1*, *MTHFD2*) upon drug treatment induced THP-1 cell cycle arrest and differentiation (PMA treatment: macrophage-like cells, VD treatment: monocyte-like cells) (24). Data indicate that serine metabolism downregulation is needed for the cell cycle arrest stage (PMA but not VD3 treatment), and that this pathway came back to normal after differentiation (PMA5Dr). VD3: Vitamin D treatment for 3 days, PMA: PMA treatment for 3 days, PMA5Dr: PMA treatment for 3 days and rest for another 5 days. Heatmap (##: q-value<0.05, FC>2; #: q-value<0.05, FC>1.5). Data are mean ± s.e.m. (n = 3 experiments). Statistical significance determined using one-way ANOVA test (*P < 0.05; **P < 0.01; and ***P < 0.001. ns: no significant difference).

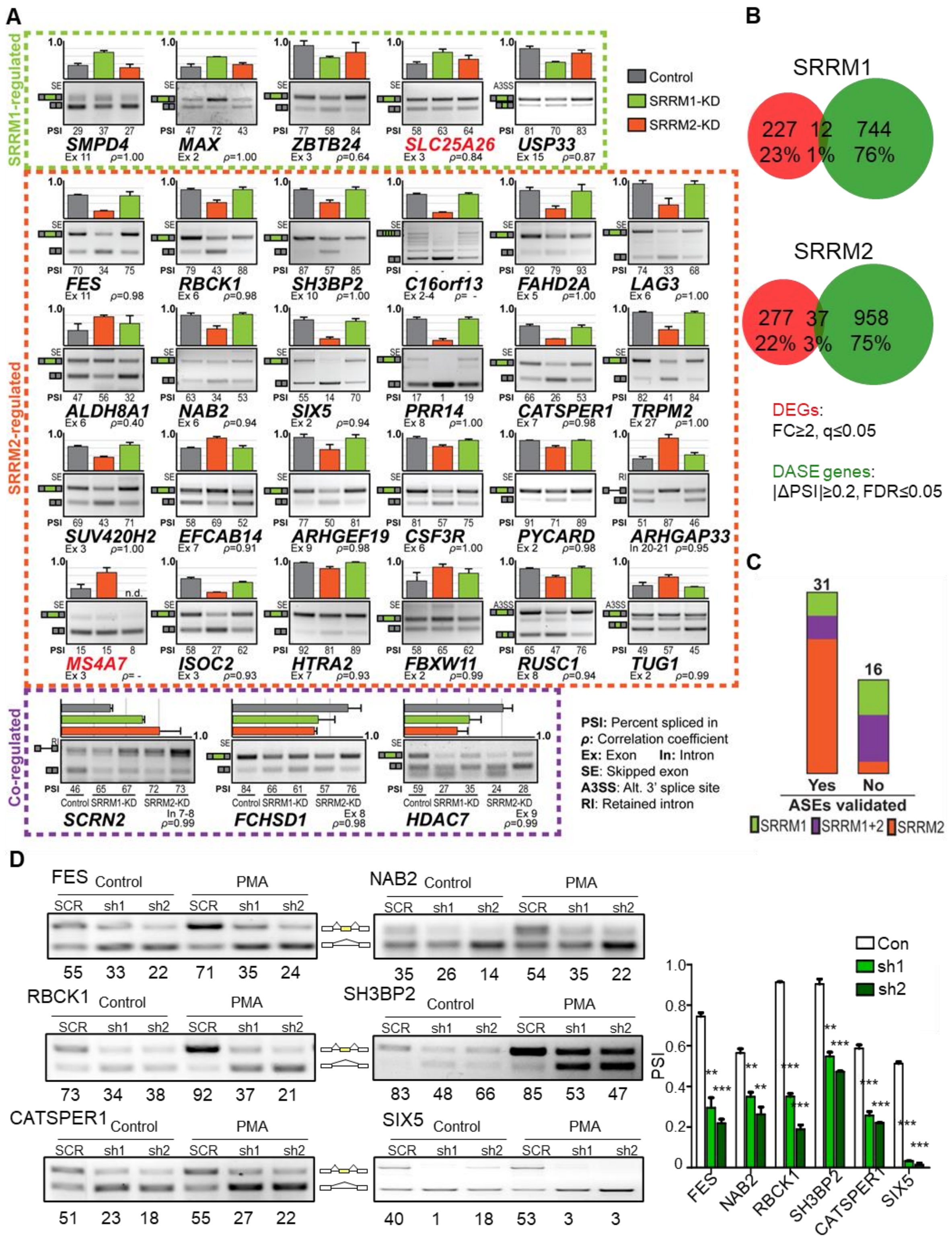

**Figure S5. DASE validation**

(**A**) Successfully validated 31 DASEs have high correlation coefficients between RT-PCR and RNA-seq data. DASEs with gene names shown in red were not validated, for reference. (**B**) Low overlap between DEG and DASE-gene datasets. (**C**) Validation of 31 out of 47 examined

DASEs using RT-PCR, sorted by SRRM1-specific, SRRM2-specific, or SRRM1/2 co-regulated. (**D**) DASE validation in THP-1 and macrophage-like THP-1 cells and statistical analysis of PSI. Data are mean ± s.e.m. (n = 3 experiments). Statistical significance determined using one-way ANOVA test (*P < 0.05; **P < 0.01; and ***P < 0.001. ns: no significant difference).

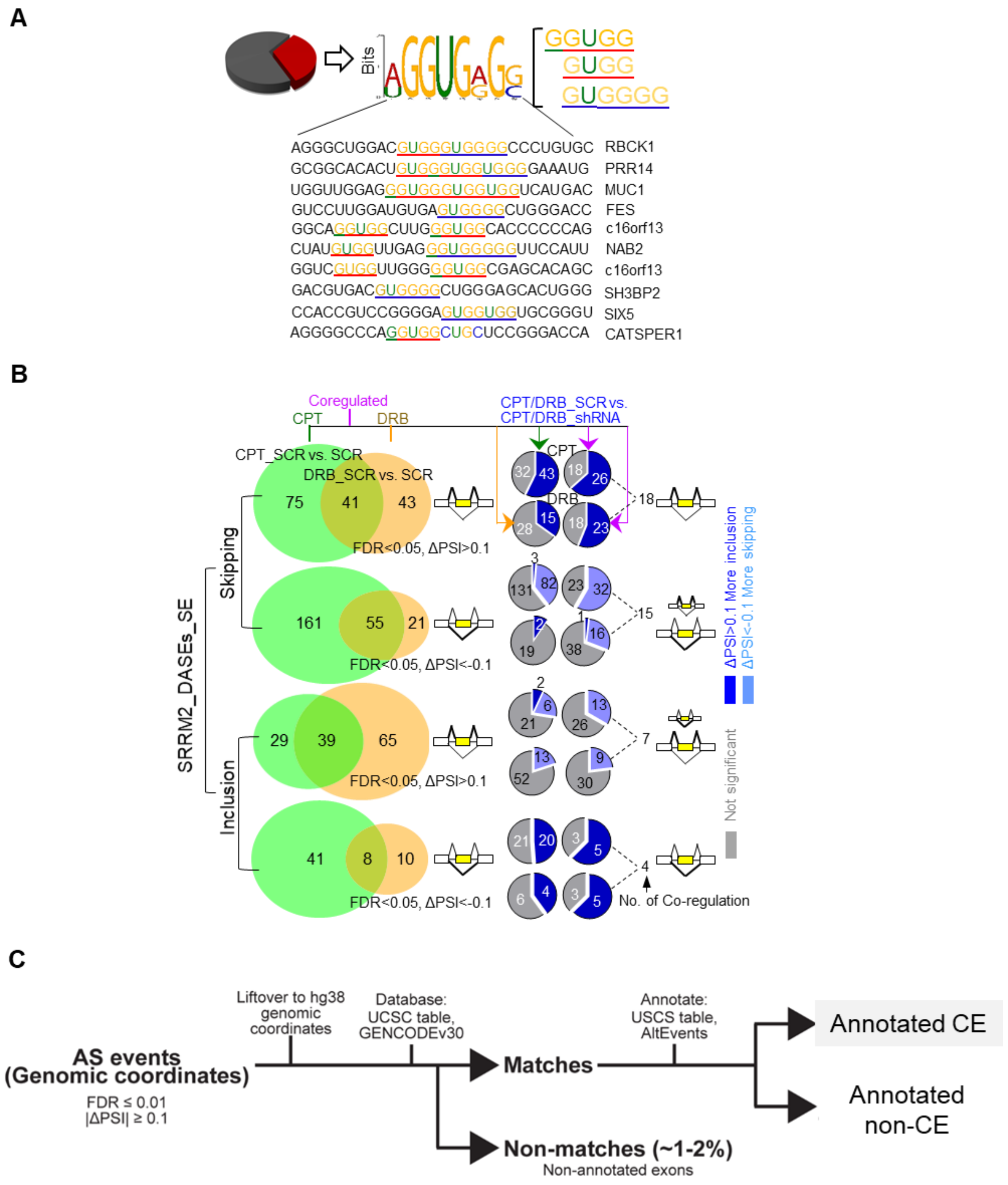

**Figure S6. Specific splicing features of SRRM2 targets**

(**A**) Motif enriched in SRRM2 regulated skipping events, as analyzed by MEME. Most of the strongly silenced splicing events had a GGUGG motif either in the intron or exon region of cassette exons. (**B**) Left: Venn diagrams showing the co-upregulation or co-downregulation of CPT and DRB on SRRM2 DASEs. Right: pie charts regulation of CPT/DRB co-regulated SRRM2 DASEs by shRNA under CPT/DRB drug treatment. Results show that, for cassette exons with increased inclusion upon DRB and/or CPT, SRRM2 knockdown is either neutral or further enhances inclusion (gray or dark blue sections in pie charts). Consistently, for cassette exons with increased skipping upon DRB and/or CPT, SRRM2 downregulation is either neutral or further enhances skipping (gray or light blue sections), with only 1-3 exceptions (dark blue sections). This result is irrespective of the effects of SRRM2 knockdown without drug, inducing either cassette exon skipping (up) or inclusion (down). (**C**) Workflow for identifying SRRM1 or SRRM2 knockdown induced annotated/non-annotated cassette exons. We converted the hg19-based genomic coordinates of the differentially spliced exons identified by rMATS to hg38-based coordinates using the UCSC LiftOver tool. We then matched them to the UCSC AltEvents table to identify annotated cassette exons, annotated non-cassette exons or completely unannotated exons.

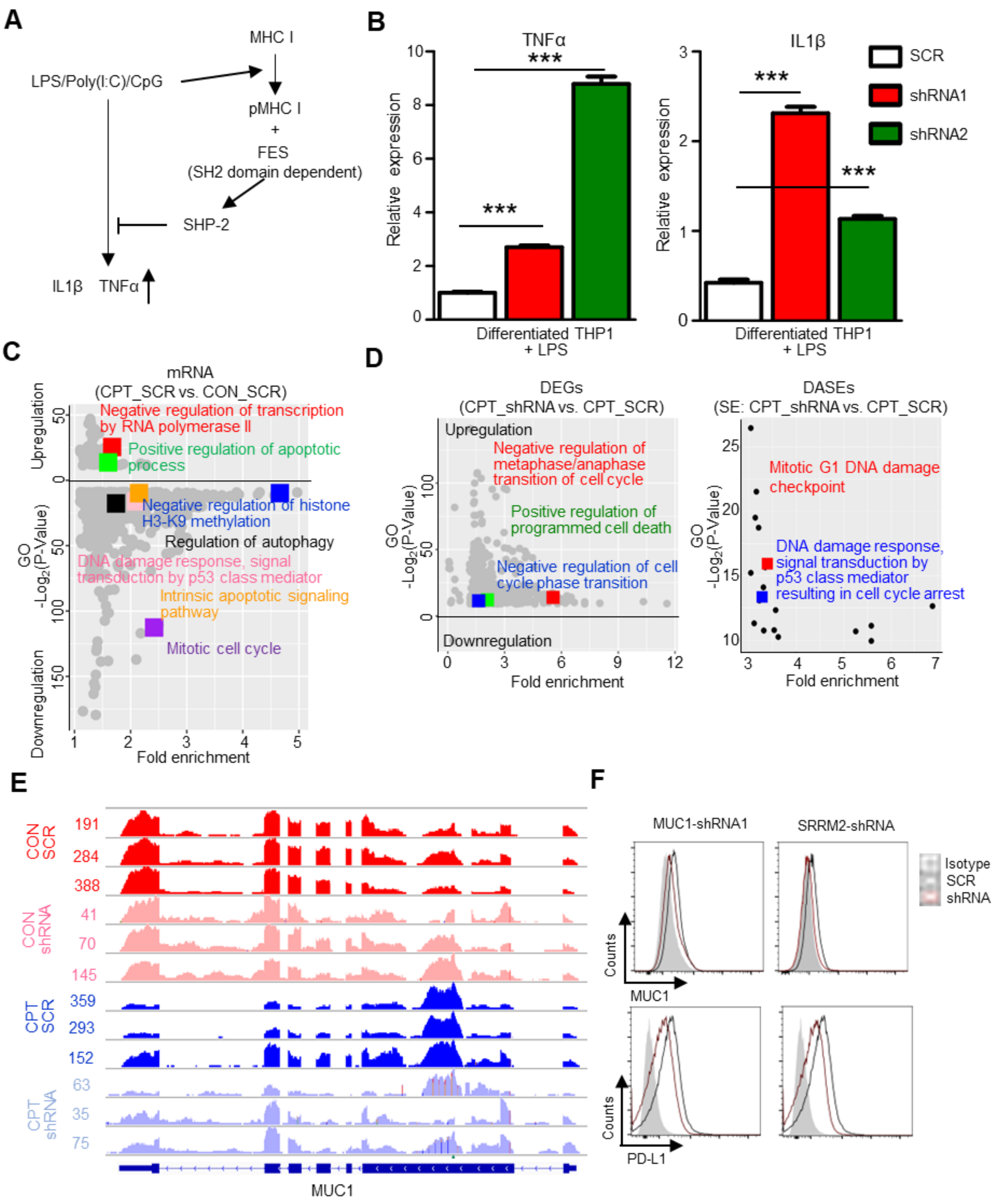

**Figure S7. SRRM2 targets have isoform-specific function in regulating innate immunity and AML cell homeostasis**

(**A**) Pathway diagram showing that FES with its SH2 domain is needed to attenuate LPS (as well as poly(I/C) and CpG)-induced proinflammatory cytokine production. (**B**) Relative expression of proinflammatory cytokines *TNFα* (left) and *IL1β* (right) in human macrophage-like THP-1 cells upon knockdown of SRRM2, under LPS (1ng/ml) for 6 h. (**C**) GO analysis revealing that CPT treatment slowed down Pol II and induced cell cycle arrest and programmed cell death. (**D**) GO analysis showing that either DEGs or DASEs of the combination treatment (SRRM2 knockdown and CPT) compared to drug (CPT) alone are enriched in cell cycle arrest and DNA damage pathways. (**E**) RNA-seq profiles to illustrate the reads and annotation of *MUC1* upon SRRM2 knockdown and CPT treatment. (**F**) FACS showing that SRRM2 knockdown significantly downregulated surface expression level of MUC1 (using MUC1 knockdown and the affected PD-L1 as a control). Data are mean ± s.e.m. (n = 3 experiments). Statistical significance determined using one-way ANOVA test (*P < 0.05; **P < 0.01; and ***P < 0.001. ns: no significant difference).

**Key Resources Table**

| Reagent or resources | Source | Identifier |
|---|---|---|
| **Antibodies** | | |
| Alexa Fluor 647 Goat anti-mouse IgG (minimal x-reactivity) Antibody | Biolegend | Cat: 405322 |
| Anti-SON antibody produced in rabbit | Sigma | Cat: HPA023535 |
| Anti-SRRM1 antibody | Abcam | Cat: ab221061 |
| Anti-SRRM2 antibody | Abcam | Cat: ab122719 |
| APC anti-human CD14 Antibody | Biolegend | Cat: 325608 (clone: HCD14) |
| DKC1 Polyclonal Antibody | Thermo Fisher/Invitrogen | Cat: PA5-28922 |
| Goat Anti-Mouse IgG H&L (HRP) | Santa Cruz | Cat: sc2005 |
| Goat Anti-Rabbit IgG H&L (HRP) | Sigma | Cat: A6154 |
| Monoclonal Anti-β-Actin antibody produced in mouse | Sigma | Cat: A5441 |
| PE anti-human CD11b | Biolegend | Cat: 982606 (clone: ICRF44) |
| PE anti-human CD227 (MUC-1) Antibody | Biolegend | Cat: 355603 |
| PE anti-human CD274 (B7-H1, PD-L1) Antibody | Biolegend | Cat: 393607 |
| Purified anti-mouse CD16/32 Antibody | Biolegend | Cat:101302 |
| SRSF2 Polyclonal Antibody | Thermo Fisher/Invitrogen | Cat: PA5-12402 |
| U2AF65 Antibody (MC3) | Santa Cruz | Cat: sc-53942 |
| Alexa Fluor 647 Goat anti-mouse IgG (minimal x-reactivity) Antibody | Biolegend | Cat: 405322 |
| Anti-SON antibody produced in rabbit | Sigma | Cat: HPA023535 |
| Anti-SRRM1 antibody | Abcam | Cat: ab221061 |
| Anti-SRRM2 antibody | Abcam | Cat: ab122719 |
| APC anti-human CD14 Antibody | Biolegend | Cat: 325608 (clone: HCD14) |
| DKC1 Polyclonal Antibody | Thermo Fisher/Invitrogen | Cat: PA5-28922 |
| Goat Anti-Mouse IgG H&L (HRP) | Santa Cruz | Cat: sc2005 |
| Goat Anti-Rabbit IgG H&L (HRP) | Sigma | Cat: A6154 |
| Monoclonal Anti-β-Actin antibody produced in mouse | Sigma | Cat: A5441 |
| PE anti-human CD11b | Biolegend | Cat: 982606 (clone: ICRF44) |
| PE anti-human CD227 (MUC-1) Antibody | Biolegend | Cat: 355603 |
| PE anti-human CD274 (B7-H1, PD-L1) Antibody | Biolegend | Cat: 393607 |
| Purified anti-mouse CD16/32 Antibody | Biolegend | Cat:101302 |
| SRSF2 Polyclonal Antibody | Thermo Fisher/Invitrogen | Cat: PA5-12402 |
| U2AF65 Antibody (MC3) | Santa Cruz | Cat: sc-53942 |

| **Reagents and Chemicals** | | |
|---|---|---|
| 5,6-Dichlorobenzimidazole 1-β-D-ribofuranoside (DRB) | Sigma | Cat: D1916 |
| 5-Aza-2′-deoxycytidine (AZA) | Sigma | Cat: A3656 |

| | | |
|---|---|---|
| Bovine Serum Albumin | Sigma | Cat: A3294 |
| Camptothecin (CPT) | Sigma | Cat: C9911 |
| DAPI (4′,6-Diamidine-2′-phenylindole dihydrochloride) | Thermo Fisher/Invitrogen | Cat: 62247 |
| Hoechst 33342 (bisBenzimide H 33342 trihydrochloride) | Thermo Fisher | Cat: H3570 |
| Leibovitz's L-15 media no phenol red | Gibco | Cat: 21083-027 |
| Phorbol 12-myristate 13-acetate (PMA) | Sigma | Cat: P8139 |
| Paraformaldehyde | Sigma | Cat: 158127 |
| Triton X-100 | Sigma | Cat: X100 |

| **Commercial Assays** | | |
|---|---|---|
| Wizard SV Genomic DNA Purification System | Promega | Cat: A2365 |
| QIAprep Spin Miniprep kit | QIAGEN | Cat: 27104 |
| QIAquick Gel Extraction kit | QIAGEN | Cat: 28706 |
| RNeasy Mini Kit | QIAGEN | Cat: 74106 |
| DreamTaq Green PCR Master Mix (2X) | Thermo Fisher | Cat: K1081 |
| μ-Dish 35 mm, high | ibidi | Cat: 81156 |
| SYBR Select Master Mix | Thermo Fisher | Cat: 4472903 |
| M-MLV Reverse Transcriptase | Promega | Cat: M170A |

| **Software and Algorithms** | | |
|---|---|---|
| Bedtools 2.25 | GitHub | https://bedtools.readthedocs.io/en/latest/ |
| Bowtie2 2.29 | Johns Hopkins University | http://bowtie-bio.sourceforge.net/bowtie2/index.shtml |
| CIDER | Pappu Lab | http://pappulab.wustl.edu/CIDER/ |
| Cufflinks 2.2.1 | Trapnell lab | http://cole-trapnell-lab.github.io/cufflinks/ |
| Fiji/ImageJ | Fiji/ImageJ | https://imagej.net/Fiji |
| FlowJo v10 | Tree Star | https://www.flowjo.com/ |
| GraphPad Prism v8 | GraphPad Software | https://www.graphpad.com/scientific-software/prism/ |
| IGV 2.9.0 | Broad Institute | http://software.broadinstitute.org/software/igv/ |
| Imaris 9.5.0 | Bitplane | https://imaris.oxinst.com/support/imaris-release-notes/9-5-0 |
| IURed2A | IURed2A | https://iupred2a.elte.hu/ |
| JavaGSEA Desktop Application | Broad Institute | http://software.broadinstitute.org/gsea |
| PLAAC | PLAAC/MIT | http://plaac.wi.mit.edu/ |
| R v3.5.2 | R-project | http://www.R-project.org/ |
| rMATS 4.0 | Xinglab/Children's Hospital of Philadelphia | http://rnaseq-mats.sourceforge.net/ |
| Samtools 1.3 | GitHub | http://www.htslib.org/ |

| | | |
|---|---|---|
| sgRNAs Design _CHOPCHOP | CHOPCHOP | https://chopchop.cbu.uib.no/ |
| TCGA | TCGA Research Network | https://www.cancer.gov/tcga |
| Tophat 2.1.1 | CCB/Johns Hopkins University | https://ccb.jhu.edu/software/tophat/index.shtml |
| ZEN lite | ZEISS | https://zeiss.com/Home/Software |

| **Deposited Data** | | |
|---|---|---|
| Bulk RNA-sequencing of SRRM2 knockdown THP-1 cells | This paper | GEO: GSE199720 |
| Bulk RNA-sequencing of CPT drug treatment AML cell line THP-1 upon SRRM2 knockdown | This paper | GEO: GSE199720 |

| **Experimental models: bacteria strains and cell lines** | | |
|---|---|---|
| E. coli BL21 (DE3) | Thermo Fisher | C600003 |
| E. coli Rosetta (DE3) | Thermo Fisher | EC0114 |
| HEK293T | ATCC | ATCC, CRL-11268 |
| THP-1 | ATCC | ATCC, TIB-202 |

| **Recombinant DNA** | | |
|---|---|---|
| pcDNA3.1(+) | Addgene | Cat: V790-20 |
| pcDNA3.1(+) -FES minigene | Addgene | Cat: V790-20 |
| pcDNA3.1(+) -SH3BP2 minigene | Addgene | Cat: V790-20 |
| pEGFP-N-NCK | This paper | N.A. |
| pEGFP-N-UPR | This paper | N.A. |
| pEGFP-N-DPR | This paper | N.A. |
| pEGFP-N-ΔNCK | This paper | N.A. |
| pEGFP-N-NCK-NES | This paper | N.A. |
| pEGFP-N-ΔUPR | This paper | N.A. |
| pEGFP-N-ΔDPR | This paper | N.A. |
| pEGFP-N-ΔUPRΔDPR | This paper | N.A. |
| pET24b(+) | Addgene | Cat: 69750-3 |
| pET24b(+) -KSR | This paper | N.A. |
| pET24b(+) -UPR3 | This paper | N.A. |
| pET24b(+) -KSR-mEGFP | This paper | N.A. |
| pET24b(+) -UPR3-mEGFP | This paper | N.A. |
| pGRIPZ-shmirRNA-SCR | This paper | N.A. |
| pGRIPZ-shmirRNA-SRRM1-1# | This paper | N.A. |
| pGRIPZ-shmirRNA-SRRM1-2# | This paper | N.A. |
| pGRIPZ-shmirRNA-SRRM2-1# | This paper | N.A. |
| pGRIPZ-shmirRNA-SRRM2-2# | This paper | N.A. |
| pLKO.1 puro | Addgene | Cat: 8453 |
| pLKO.1 SCR | This paper | N.A. |
| pLKO-shRNA-FES | This paper | N.A. |
| pLKO-shRNA-FES-iso1/2 | This paper | N.A. |
| pLKO-shRNA-SRRM1-1# | This paper | N.A. |
| pLKO-shRNA-SRRM1-2# | This paper | N.A. |
| pLKO-shRNA-SRRM1-3# | This paper | N.A. |
| pLKO-shRNA-SRRM2-1# | This paper | N.A. |

| | | |
|---|---|---|
| pLKO-shRNA-SRRM2-2# | This paper | N.A. |
| pLKO-shRNA-SRRM2-3# | This paper | N.A. |
| pLKO-shRNA-MUC1-1# | This paper | N.A. |
| pLKO-shRNA-MUC1-2# | This paper | N.A. |
| pmCherry-N1 | This paper | N.A. |
| pmEGFP-N | This paper | N.A. |
| pMD2.G | Addgene | Cat: 12259 |
| Other SRRM2 mutants for domain mapping | This paper | N.A. |
| psPAX2 | Addgene | Cat: 12260 |
| pUC19 | Addgene | Cat: 50005 |
| pX330 | Addgene | Cat: 42230 |
| pX330-sgRNA-SRRM2 | This paper | N.A. |
| pX330-sgRNA-SRSF2 | This paper | N.A. |
| pX459 | Addgene | Cat: 62988 |
| pET24b(+) | Addgene | Cat: 69750-3 |
| pET24b(+) -KSR | This paper | N.A. |
| pET24b(+) -UPR3 | This paper | N.A. |
| pET24b(+) -KSR-mEGFP | This paper | N.A. |
| pET24b(+) -UPR3-mEGFP | This paper | N.A. |
| pGRIPZ-shmirRNA-SCR | This paper | N.A. |
| pGRIPZ-shmirRNA-SRRM1-1# | This paper | N.A. |
| pGRIPZ-shmirRNA-SRRM1-2# | This paper | N.A. |
| pGRIPZ-shmirRNA-SRRM2-1# | This paper | N.A. |
| pGRIPZ-shmirRNA-SRRM2-2# | This paper | N.A. |
| pLKO.1 puro | Addgene | Cat: 8453 |
| pLKO.1 SCR | This paper | N.A. |
| pLKO-shRNA-FES | This paper | N.A. |
| pLKO-shRNA-FES-iso1/2 | This paper | N.A. |
| pLKO-shRNA-SRRM1-1# | This paper | N.A. |
| pLKO-shRNA-SRRM1-2# | This paper | N.A. |
| pLKO-shRNA-SRRM1-3# | This paper | N.A. |
| pLKO-shRNA-SRRM2-1# | This paper | N.A. |
| pLKO-shRNA-SRRM2-2# | This paper | N.A. |
| pLKO-shRNA-SRRM2-3# | This paper | N.A. |
| pLKO-shRNA-MUC1-1# | This paper | N.A. |
| pLKO-shRNA-MUC1-2# | This paper | N.A. |
| pmCherry-N1 | This paper | N.A. |
| pmEGFP-N | This paper | N.A. |
| pMD2.G | Addgene | Cat: 12259 |
| Other SRRM2 mutants for domain mapping | This paper | N.A. |
| psPAX2 | Addgene | Cat: 12260 |
| pUC19 | Addgene | Cat: 50005 |
| pX330 | Addgene | Cat: 42230 |
| pX330-sgRNA-SRRM2 | This paper | N.A. |
| pX330-sgRNA-SRSF2 | This paper | N.A. |
| pX459 | Addgene | Cat: 62988 |

**Supplementary Table 1**

***Oligos and peptides***

| Primers for AS Validation | | |
|---|---|---|
| **Gene** | **F/R** | **Sequence (5'-3')** |
| ALDH8A1 | F | GACGCCAACCTGGATGAGTG |
| | R | CACCGTGGGAAGCATAAAGTAGC |
| ARHGAP33 | F | ACCCTACTTACCAAGGCAAC |
| | R | CCGATCTCATAGTACAGGTTCTC |
| ARHGEF19 | F | ACCTGCCCTATGTCACCAAC |
| | R | TGTCTTCGTCTTCAGAGCC |
| ARMC10 | F | CTCGGGATACGCTCTTCG |
| | R | TTGACTCCAGCAGGTAAAGG |
| c16orf13 | F | CTGCGGCAGTACCTGGATC |
| | R | GGCATGTCCACCATCCTCTC |
| CATSPER1 | F | ACCTTGCTCACGCTGGATG |
| | R | CTCCTGCTTCGCTTTCTCAAG |
| CSF3R | F | GACCTGAGACCCACCTACC |
| | R | GCTCCAGTCGCTCCAGT |
| EFCAB14 | F | GCGTCCATCTTGCTGTG |
| | R | ATCTGTTCTGTTGGTGACTTGA |
| FAHD2A | F | GGAAAGAAAGGCAAGCACATCAAG |
| | R | GGGAGACCCAGGCTATCAG |
| FBXW11 | F | TCGGTGATTGAGGACAAGAC |
| | R | CATTCGTGAAATAAGATGTTCCACAA |
| FCHSD1 | F | ATCTTCCACTCTCGGACCA |
| | R | GACCTCTGCGGCTTCCA |
| FES | F | GCCACATCTCAGGAATCTTC |
| | R | GCAGGACAACACCACTCTTC |
| HDAC7 | F | CTCCTCCCCAAGTAGTAGC |
| | R | GGTCCTGCGGTCACTGT |
| HTRA2 | F | ATTGGGGTGATGATGCTGA |
| | R | CAACTGGGATTGGGTTCG |
| ISOC | F | TCCTCTGTCCTGTTCCTGT |
| | R | GGCGTCCACCACCACAT |
| LAG3 | F | CACTGTCACATTGGCAATCATCAC |
| | R | GGTTCTTGCTCCAGCTCCTC |
| MAX | F | CTCCCCTTCACCCCTTC |
| | R | ATTATGATGAGCCCGTTTGTC |
| MUC1 | F1 | TGGAAGATCCCAGCACCGA |
| | F2 | ATGGCACATCACTCACATTTCAGA |
| | R | GGCACATCACTCACTGAACTTAATATTG |
| NAB2 | F | GAGAACAGAGTCACCCTGAAATC |
| | R | CACTTTCACGCTGCTCCTG |
| PRR14 | F | AGCTGAGCCCATGAGGATAGTTC |
| | R | GGTTCTGAAAGCACCTGGTCC |
| PYCARD | F | TGGACCTCACCGACAAG |
| | R | TGCCTGGTACTGCTCATC |
| RBCK1 | F | ACCTTCATCAACAAGCCCAC |
| | R | GCACGAGTAGGTGTTGTCAATG |
| RUSC1 | F | GCAGTTGGAGCTGTGGTTTTC |
| | R | GACTTTAGGGCTCCATTCTCATTTTC |

| | | |
|---|---|---|
| SCRN2 | F | GTGTTCAAACCTTTCATCTTCGGGA |
| | R | TGTTTCTGCTGGAGCTGCTG |
| SH3BP2 | F | GATGGGCAGAGTTTCAGGAG |
| | R | ATGCAGTAGAGTCCATCCTG |
| SMPD4 | F | TGGCTTCATCACTATTCCTTG |
| | R | CTGGCTTCAGGCTGTTG |
| SIX5 | F | CGAGGAGACAGTCTACTGCTTC |
| | R | CCGTCTCTGGCTTCAGTG |
| SUV420H2 | F | GACAGAGTGACAGCACGAGAAC |
| | R | GCACGGAGGTAGCGATAGAC |
| USP33 | F | GCATTCATCAAGTCATCCAAC |
| | R | AAGAAGGCAGCAAGACAATC |
| TRPM2 | F | TCTATGACCCACCCTTTTACAC |
| | R | ATCCTCGTTCCGCCTCCAC |
| TUG1 | F | ACTGTTGACCTTGCTGTGAG |
| | R | ATCTCGGGCAAAAATCCAGG |
| ZBTB24 | F | GCTGTAAAGACTGTGGCAAGG |
| | R | TGTATGAACTCGGTAATGGCTCT |

| qPCR primers | | |
|---|---|---|
| **Gene** | **F/R** | **Sequence (5'-3')** |
| Actin | F | CCAGAGGCGTACAGGGATAG |
| | R | CCAACCGCGAGAAGATGA |
| ATF4 | F | TCAGTCCCTCCAACAACAGCAAG |
| | R | CAACGTGGTCAGAAGGTCATC |
| CBS | F | TGGTGGACAAGTGGTTCAAGAG |
| | R | GCACTGAGTCGGGCAGAATG |
| CD11b | F | CAGGGAAGTGGCAAGGAATG |
| | R | GAGCCAGGTCATAAGTCACAAC |
| CD14 | F | GTTCGGAAGACTTATCGACCATG |
| | R | GCTGAGGTTCGGAGAAGTTG |
| CD163 | F | CAAGAACTGGCAATGGGGTGG |
| | R | TGCTTCACTTCAACACGTCCAG |
| CD36 | F | ACAGGCACAGAAGTTTACAGACAG |
| | R | CTGTGTTGTCCTCAGCGTCC |
| MTHFD2 | F | AGGACGAATGTGTTTGGATCAG |
| | R | TTCCAGCCACAACCACATTC |
| IFNB1 | F | GAGCTACAACTTGCTTGGATTCC |
| | R | ATCTCCTCAGGGATGTCAAAGTTC |
| IL1B | F | ATGGACAAGCTGAGGAAGATGC |
| | R | CAGTTCAGTGATCGTACAGGTGC |
| PHGDH | F | GGGAACAGAGCTGAATGGAAAG |
| | R | CTGCTGAACACCAAAGGAGG |
| PSAT1 | F | CGTGGTGATTGTCCGTGATG |
| | R | TCCAGAACCAAGCCCATGAC |
| SHMT2 | F | CATCGTCACCACCACTACTCAC |
| | R | ACGGCAAAGTTGATTCGGTC |
| SRRM1 | F | CAGAGACGCCAATACAGACGAC |
| | R | GCTTTCGTGGTGATGGAGAAG |
| SRRM2 | F | TGAACCAGGTACTACCAGCACAC |
| | R | GGAGTGGGAGCAGGTGGTTC |

| shRNA |
|---|

| Gene | shRNA | shRNA targeted sequence |
|---|---|---|
| SCR (Scramble) | 1# | CCCTAAGGTTAAGTCGCCCTC |
| | 2# | CAACAAGATGAAGAGCACCAC |
| FES | Target all isoforms | GCTGAGATCACCAGCCAAACT |
| | Target longer isoform (iso 1 and 2) | GCTGCAGCTCATTCCGGAGGT |
| SRRM1 | 1# | GAACTCCGCCAAGAAGAATGC |
| | 2# | GCATCCTTGTCTGGGAGTAGC |
| | 3# | GGTTATCTCCTTCAGCAAGTC |
| SRRM2 | 1# | GCATCCAGATCTCCAATAAGA |
| | 2# | GCCTGAGTCTTCACTGGTATT |
| | 3# | GCTCAACAACAAGTGTCTTAC |
| MUC1 | 1# | GCCTCTCCAATATTAAGTTCA |
| | 2# | GCCAGGATCTGTGGTGGTACA |

| shmirRNA | | |
|---|---|---|
| Gene | shmirRNA | shmirRNA sequence |
| miR30-SCR | 1# | TGCTGTTGACAGTGAGCGACCTAAGGTTAAGTCGCCCT CCTAGTGAAGCCACAGATGTAGGAGGGCGACTTAACCT TAGGCTGCCTACTGCCTCGGA |
| SRRM1 | 1# | TGCTGTTGACAGTGAGCGAAACTCCGCCAAGAAGAATG CCTAGTGAAGCCACAGATGTAGGCATTCTTCTTGGCGGA GTTCTGCCTACTGCCTCGGA |
| | 2# | TGCTGTTGACAGTGAGCGACATCCTTGTCTGGGAGTAG CTTAGTGAAGCCACAGATGTAAGCTACTCCCAGACAAGG ATGCTGCCTACTGCCTCGGA |
| SRRM2 | 1# | TGCTGTTGACAGTGAGCGACATCCAGATCTCCAATAAGA CTAGTGAAGCCACAGATGTAGTCTTATTGGAGATCTGGA TGCTGCCTACTGCCTCGGA |
| | 2# | TGCTGTTGACAGTGAGCGACCTGAGTCTTCACTGGTATT CTAGTGAAGCCACAGATGTAGAATACCAGTGAAGACTCA GGCTGCCTACTGCCTCGGA |

| Minigene Primers | | |
|---|---|---|
| **Gene** | **F/R** | **Primers** |
| FES | F | CGGGATCCTCAGGAGCAGGAGCGAGAG |
| | R | CCCTCGAGCTGACCCACACTCCCCAAG |
| SH3BP2 | F | GAAGATCTTGCCCCTCCAGGCGATCA |
| | R | CCCTCGAGACCTTCCCCGACTTGGTAGAGG |

| Peptide | | |
|---|---|---|
| **Peptide** | **Method** | **Sequence** |
| K(SR) | SEC | KQKKKKKKKDRGRRSESSSPRRERKKSSKKKKHRSESES KKRKHRSPTPKSKRKSKDKKRK |
| $UPR_3$ | SEC | GRSRSRTPARRGRSRSRTPARRGRSRSRTPARR |

| sgRNA | | |
|---|---|---|
| **Target** | **type** | **Sequence** |
| SRRM2 | Knock-in | AGACCTGCAAGAGAAGATATGGG |
| SRSF2 | Knock-in | GACATTACCATTTTCTTAAGAGG |

### Supplementary Table 2

#### *AS Validation*

| Gene | Band | Band size | Exon (Ex) | Event | Δbp | PSI (RNA-Seq) | | | ΔPSI | | PSI (RT-PCR) | | | ΔPSI | | ρ |
|---|---|---|---|---|---|---|---|---|---|---|---|---|---|---|---|---|
| | | | | | | SCR | SRRM1 | SRRM2 | SRRM1 | SRRM2 | SCR | SRRM1 | SRRM2 | SRRM1 | SRRM2 | |
| FES | Upper | 351 | 10 | SE Ex11 | 210 | 0.618 | 0.583 | 0.196 | 0.034 | 0.421 | 0.699 | 0.749 | 0.344 | -0.050 | 0.355 | 0.983 |
| | lower | 141 | 12 | | | 0.015 | 0.104 | 0.014 | | | | | | | | |
| RBCK1 | Upper | 390 | 5 | SE Ex6 | 161 | 0.761 | 0.748 | 0.424 | 0.013 | 0.337 | 0.786 | 0.876 | 0.426 | -0.090 | 0.360 | 0.975 |
| | lower | 229 | 8 | | | 0.029 | 0.065 | 0.068 | | | | | | | | |
| SH3BP2 | Upper | 206 | 9 | SE Ex10 | 56 | 0.922 | 0.949 | 0.520 | -0.027 | 0.402 | 0.865 | 0.846 | 0.567 | 0.019 | 0.298 | 0.994 |
| | lower | 150 | 11 | | | 0.003 | 0.007 | 0.168 | | | | | | | | |
| RUSC1 | Upper | 591 | 7 | A3'SS Ex8 | 318 | 0.785 | 0.799 | 0.555 | -0.014 | 0.230 | 0.647 | 0.764 | 0.466 | -0.117 | 0.181 | 0.939 |
| | lower | 273 | 8 | | | 0.030 | 0.016 | 0.047 | | | | | | | | |
| c16orf13 | Upper | 527 | 1 | SE Ex2,3,4 | 291 | 0.634 | 0.632 | 0.166 | 0.002 | 0.469 | 0.648 | 0.661 | 0.300 | -0.013 | 0.348 | 0.999 |
| | lower | 236 | 5+6 | | | 0.024 | 0.034 | 0.017 | | | | | | | | |
| FAHD2A | Upper | 296 | 4 | SE Ex5 | 163 | 0.747 | 0.707 | 0.258 | 0.040 | 0.489 | 0.924 | 0.929 | 0.788 | -0.004 | 0.136 | 0.995 |
| | lower | 133 | 6 | | | 0.083 | 0.178 | 0.071 | | | | | | | | |
| TUG1 | Upper | 243 | 1 | A3'SS Ex2 | 69 | 0.343 | 0.293 | 0.560 | 0.050 | -0.217 | 0.488 | 0.449 | 0.574 | 0.039 | -0.086 | 0.991 |
| | lower | 174 | 2 | | | 0.083 | 0.009 | 0.057 | | | | | | | | |
| LAG3 | Upper | 486 | 5 | SE Ex6 | 243 | 0.924 | 0.902 | 0.364 | 0.023 | 0.561 | 0.738 | 0.677 | 0.334 | 0.060 | 0.404 | 0.995 |
| | lower | 243 | 8 | | | 0.076 | 0.085 | 0.172 | | | | | | | | |
| SCRN2 | Upper | 467 | 7 | IR In7-8 | 260 | 0.373 | 0.619 | 0.749 | -0.246 | -0.376 | 0.458 | 0.662 | 0.722 | -0.204 | -0.264 | 0.992 |
| | lower | 207 | 8 | | | 0.017 | 0.016 | 0.157 | | | | 0.650 | 0.717 | | | |
| | | | | | | | | | | | | 0.673 | 0.726 | | | |
| ARHGAP33 | Upper | 369 | 20 | IR In20-21 | 84 | 0.316 | 0.427 | 0.857 | -0.111 | -0.541 | 0.510 | 0.461 | 0.868 | 0.049 | -0.358 | 0.954 |
| | lower | 285 | 21 | | | 0.056 | 0.058 | 0.094 | | | | | | | | |
| ALDH8A1 | Upper | 339 | 5 | SE Ex6 | 162 | 0.401 | 0.582 | 0.763 | -0.181 | -0.362 | 0.464 | 0.315 | 0.563 | 0.149 | -0.099 | 0.398 |
| | lower | 177 | 7 | | | 0.175 | 0.210 | 0.041 | | | | | | | | |
| NAB2 | Upper | 407 | 5 | SE Ex6 | 192 | 0.809 | 0.815 | 0.445 | -0.006 | 0.364 | 0.632 | 0.535 | 0.341 | 0.097 | 0.291 | 0.940 |
| | lower | 215 | 7 | | | 0.053 | 0.042 | 0.081 | | | | | | | | |
| SIX5 | Upper | 1172 | 1 | SE Ex2 | 806 | 0.680 | 0.630 | 0.184 | 0.050 | 0.495 | 0.549 | 0.705 | 0.142 | -0.155 | 0.407 | 0.935 |
| | lower | 366 | 3 | | | 0.068 | 0.066 | 0.046 | | | | | | | | |
| PRR14 | Upper | 756 | 7 | SE Ex8 | 586 | 0.704 | 0.690 | 0.139 | 0.014 | 0.565 | 0.169 | 0.191 | 0.013 | -0.022 | 0.156 | 0.991 |
| | lower | 170 | 9 | | | 0.050 | 0.052 | 0.046 | | | | | | | | |
| CATSPER1 | Upper | 192 | 6 | SE Ex7 | 64 | 0.519 | 0.480 | 0.183 | 0.039 | 0.335 | 0.657 | 0.529 | 0.256 | 0.128 | 0.402 | 0.978 |
| | lower | 128 | 8 | | | 0.044 | 0.063 | 0.008 | | | | | | | | |
| TRPM2 | Upper | 272 | 26 | SE Ex27 | 102 | 0.882 | 0.856 | 0.424 | 0.026 | 0.458 | 0.824 | 0.837 | 0.412 | -0.013 | 0.412 | 0.997 |
| | lower | 170 | 29 | | | 0.021 | 0.038 | 0.085 | | | | | | | | |
| SUV420H2 | Upper | 287 | 2 | SE Ex3 | 166 | 0.642 | 0.683 | 0.358 | -0.041 | 0.284 | 0.688 | 0.713 | 0.429 | -0.025 | 0.259 | 0.999 |
| | lower | 122 | 4 | | | 0.028 | 0.065 | 0.025 | | | | | | | | |
| FBXW11 | Upper | 246 | 1 | SE Ex2 | 63 | 0.456 | 0.669 | 0.814 | -0.213 | -0.358 | 0.582 | 0.617 | 0.653 | -0.035 | -0.071 | 0.993 |
| | lower | 183 | 3 | | | 0.219 | 0.116 | 0.068 | | | | | | | | |
| EFCAB14 | Upper | 440 | 5 | SE Ex7 | 192 | 0.585 | 0.595 | 0.856 | -0.010 | -0.271 | 0.584 | 0.518 | 0.686 | 0.066 | -0.102 | 0.909 |
| | lower | 248 | 8 | | | 0.026 | 0.055 | 0.036 | | | | | | | | |
| ARHGEF19 | Upper | 228 | 8 | SE Ex9 | 130 | 0.878 | 0.859 | 0.555 | 0.019 | 0.323 | 0.767 | 0.807 | 0.504 | -0.040 | 0.263 | 0.985 |
| | lower | 98 | 10 | | | 0.078 | 0.093 | 0.130 | | | | | | | | |
| CSF3R | Upper | 527 | 5 | SE Ex6 | 188 | 0.851 | 0.805 | 0.644 | 0.046 | 0.208 | 0.805 | 0.748 | 0.570 | 0.057 | 0.235 | 1.000 |
| | lower | 239 | 8 | | | 0.036 | 0.034 | 0.040 | | | | | | | | |
| PYCARD | Upper | 296 | 1 | SE Ex2 | 57 | 0.795 | 0.809 | 0.619 | -0.014 | 0.176 | 0.912 | 0.888 | 0.709 | 0.024 | 0.203 | 0.985 |
| | lower | 239 | 3 | | | 0.008 | 0.015 | 0.040 | | | | | | | | |
| ISOC | Upper | 363 | 2 | SE Ex3 | 210 | 0.526 | 0.430 | 0.158 | 0.096 | 0.368 | 0.578 | 0.622 | 0.269 | -0.043 | 0.309 | 0.933 |
| | lower | 153 | 4 | | | 0.032 | 0.024 | 0.005 | | | | | | | | |
| HTRA2 | Upper | 228 | 6 | SE Ex7 | 96 | 0.930 | 0.944 | 0.773 | -0.014 | 0.157 | 0.921 | 0.887 | 0.808 | 0.034 | 0.113 | 0.931 |
| | lower | 132 | 8 | | | 0.011 | 0.017 | 0.053 | | | | | | | | |
| SMPD4 | Upper | 232 | 10 | SE Ex11 | 87 | 0.368 | 0.704 | 0.300 | -0.336 | 0.067 | 0.290 | 0.372 | 0.266 | -0.082 | 0.025 | 0.998 |
| | lower | 145 | 12 | | | 0.052 | 0.047 | 0.087 | | | | | | | | |
| USP33 | Upper | 177 | 14 | A3SS Ex15 | 24 | 0.793 | 0.445 | 0.679 | 0.347 | 0.113 | 0.808 | 0.695 | 0.835 | 0.113 | -0.027 | 0.873 |
| | lower | 153 | 15 | | | 0.059 | 0.019 | 0.079 | | | | | | | | |
| MAX | Upper | 149 | 1 | SE Ex2 | 27 | 0.313 | 0.584 | 0.379 | -0.271 | -0.066 | 0.469 | 0.716 | 0.434 | -0.246 | 0.036 | 0.939 |
| | lower | 122 | 3 | | | 0.073 | 0.051 | 0.007 | | | | | | | | |
| ZBTB24 | Upper | 320 | 2 | SE Ex3 | 168 | 0.880 | 0.558 | 0.697 | 0.322 | 0.183 | 0.767 | 0.577 | 0.838 | 0.190 | -0.071 | 0.649 |
| | lower | 152 | 4 | | | 0.129 | 0.026 | 0.270 | | | | | | | | |
| HDAC7 | Upper | 232 | 8 | SE Ex9 | 111 | 0.754 | 0.498 | 0.401 | 0.256 | 0.352 | 0.586 | 0.311 | 0.259 | 0.275 | 0.327 | 0.993 |
| | lower | 112 | 10 | | | 0.075 | 0.092 | 0.082 | | | | 0.272 | 0.240 | | | |
| | | | | | | | | | | | | 0.350 | 0.279 | | | |
| ARMC10 | Upper | 212 | 1 | SE Ex2 | 105 | 0.302 | 0.529 | 0.390 | -0.226 | -0.088 | 0.396 | 0.503 | 0.452 | -0.107 | -0.056 | 0.988 |
| | lower | 107 | 3 | | | 0.088 | 0.042 | 0.055 | | | | 0.453 | 0.435 | | | |
| | | | | | | | | | | | | 0.553 | 0.469 | | | |
| FCHSD1 | Upper | 252 | 7 | SE Ex8 | 129 | 0.882 | 0.652 | 0.625 | 0.230 | 0.257 | 0.887 | 0.694 | 0.726 | 0.193 | 0.160 | 0.968 |
| | lower | 123 | 9 | | | 0.103 | 0.018 | 0.133 | | | | 0.707 | 0.639 | | | |
| | | | | | | | | | | | | 0.681 | 0.814 | | | |

**Supplementary Video Legends**

**Video 1**. Live-cell images of two fusion events among three condensates of knock-in EGFP-SRRM2 in HEK293T cells.

**Video 2**. Representative 4D (real-time 3D) live-cell imaging (maximum intensity projection of stacks) for tracking showing the dynamic distribution of SRRM2-EGFP and SRSF2-mCherry from cytokinesis to interphase in HEK293T.

**Video 3**. Time-lapse of $UPR_3$ droplets at concentration of 20 μM under 150 mM NaCl.
